# A cross-species single-cell epigenome kidney atlas identifies epithelial cells as a driver of epigenetic aging

**DOI:** 10.64898/2026.01.22.700871

**Authors:** Hyeonsoo Jeong, Blue B. Lake, Dinh Diep, Xuwen Li, Qi Yan, Debora L. Gisch, Stephanie Reinert, Michael T. Eadon, Joseph P. Gaut, Sanjay Jain, Kun Zhang

## Abstract

Epigenetic aging is a hallmark of chronic diseases, arising from sustained injuries and unresolved repairs. To investigate cell-type-specific epigenetic alterations, we built a cross-species single-cell multi-omics atlas of DNA methylomes, chromatin accessibilities, and transcriptomes on healthy, injured (human) and aging (mouse) kidneys. We identified accelerated epigenetic aging dominated by tubular epithelia in diseased kidneys. The pathological state mirrors transcriptional trajectories observed in normal aging, driven by the preferential dysregulation of lineage-specific genes lacking CpG islands. Spatially, these epigenetic changes mapped to pathological niches of failed repair. Co-profiling of single-cell DNA methylation and 3D genome architecture revealed that epithelial repair states in disease undergo significant higher-order genome reorganizations, activating genes associated with renal decline. Our findings demonstrate that epithelial aging is driven by a collapse of 3D chromatin structure and local methylome integrity, which silences cell identity and promotes a non-resolving repair state.

## Main

Chronic diseases, such as chronic kidney disease (CKD), represent a significant public health challenge, placing an immense burden on patients and healthcare systems worldwide ^1,2^. CKD often stems from progressive decline in kidney function due to prolonged injury and inflammation, leading to impaired functional repair of nephron cell types. Instead of resolving, some injured cells can adopt a persistent, dysfunctional state that contributes to fibrosis and organ failure ^3–6^. Epigenetic mechanisms are essential for establishing and maintaining cell identity and modulating cellular responses to stress. While these mechanisms normally guide successful repair, epigenetic dysregulation can shift tissues from adaptive repair to a persistent pathological state ^7,8^. However, with multiple cell types present in the kidney, the cell type-specific epigenetic programs that drive this failure in chronic diseases are not yet well understood.

Dissecting these mechanisms of CKD pathogenesis has been challenging due to the complex cellular heterogeneity of the kidney. While recent single-cell studies have provided insights into transcriptomes and chromatin accessibilities ^5,9–11^, a cell type-resolved view of the deeper epigenetic layers governing cell identity remains missing. Specifically, DNA methylation provides a stable, long-term record of cellular states ^12,13^, while 3D genome architecture governs the physical topology required for gene regulation ^14,15^. Capturing these layers is essential to understand how regulatory circuits are rewired in chronic conditions. Furthermore, while aging is known to be a primary risk factor for CKD ^16^, the interplay between normal age-related epigenetic drift and the specific alterations driving disease pathology remains to be elucidated.

To address these gaps, we built a cross-species, multimodal kidney single-cell epigenome atlas spanning human health and CKD, alongside with young and aged mouse kidneys. This cross-species data set has enabled an examination of conserved mechanisms of aging and chronic disease. By generating single-cell DNA methylomes with matched single-cell multiomes (RNA/ATAC) from the same donors, we identified the cell-type-specific epigenetic changes associated with aging and disease, highlighting the tubular epithelium as a primary driver of accelerated epigenetic aging. Our atlas uniquely integrates these modalities with spatial transcriptomics (Xenium) and single-cell methyl-Hi-C (scMethyl-Hi-C ^17^) to map these changes to specific tissue niches and revealed a collapse of higher-order chromatin organization. Our findings provide a key resource and insights into spatially anchored gene expression changes integrated with single cell epigenetic changes across age span with cross-species validations.

## Results

### A multimodal epigenome atlas of the human and mouse kidneys

We applied a cellular combinatorial indexing method (sciMET ^18^) to generate a cross-species single-cell DNA methylation atlas comprising 64,203 high-quality nuclei. This dataset spans 12 human donors (34,456 nuclei from 7 CKD patients and 5 age-matched healthy controls; Supplementary Table 1) and 6 mice (29,747 nuclei from 3 young and 3 aged animals; Supplementary Table 2), yielding a median of 0.91 million CpG sites (approximately 3.1% CpG coverage of the human genome) per cell. To minimize technical variation, human samples were pooled for processing in one batch and deconvoluted based on SNPs. Donor-specific variants from whole-genome DNA sequencing validated multiplexing accuracy, with 99.2% of sciMET nuclei assigned to the correct donor of origin.

To link these epigenetic profiles to gene regulation, tissue niches, and chromatin architecture, we integrated the single-cell DNA methylation atlas with matched multimodal datasets. We generated single-cell multiomes profiles for the matched human (142,459 nuclei from 9 samples) and mouse samples (36,563 nuclei from 6 samples) to directly connect DNA methylation with chromatin accessibility and gene expression (Supplementary Table 1). We further contextualized these findings by generating spatial transcriptomics maps using Xenium for representative healthy, acute kidney injury (AKI), and CKD samples (383,771 cells from 4 samples), and defined the baseline 3D chromatin compartments using scMethyl-Hi-C on a healthy reference donor. This multi-layered approach establishes a rigorous framework for characterizing cell-type-specific epigenetic drivers of aging and disease.

We first sought to accurately identify and annotate kidney cell types based on their unique epigenetic profiles. Unsupervised clustering of human single-cell methylomes successfully identified all known major kidney cell types (Fig. 1a). We annotated these clusters based on promoter hypomethylation of canonical marker genes, such as PODXL for podocytes (POD) and SLC34A1 for proximal tubule (PT) cells (Supplementary Fig. 1). Assessment of orthologous genes ensured consistent identification of matched cell types in the mouse kidney. Global DNA methylation levels were largely stable, ranging from 69.1% to 81.7% across most lineages, with the exception of immune cells, which exhibited greater heterogeneity of CpG methylation consistent with their diverse functional states (Fig. 1b). Aggregating single-cell profiles yielded pseudo-bulk cell type methylomes ranging from 28.35 to 29.2 million CpGs, ensuring robust detection of regulatory elements. Validation against reference bulk methylomes ^19^ confirmed the high accuracy of our cell type assignments, while our single-cell approach provided substantially higher resolution, enabling the distinction of previously unresolved subpopulations within the tubular epithelium (Supplementary Fig. 2).

**Fig. 1.**
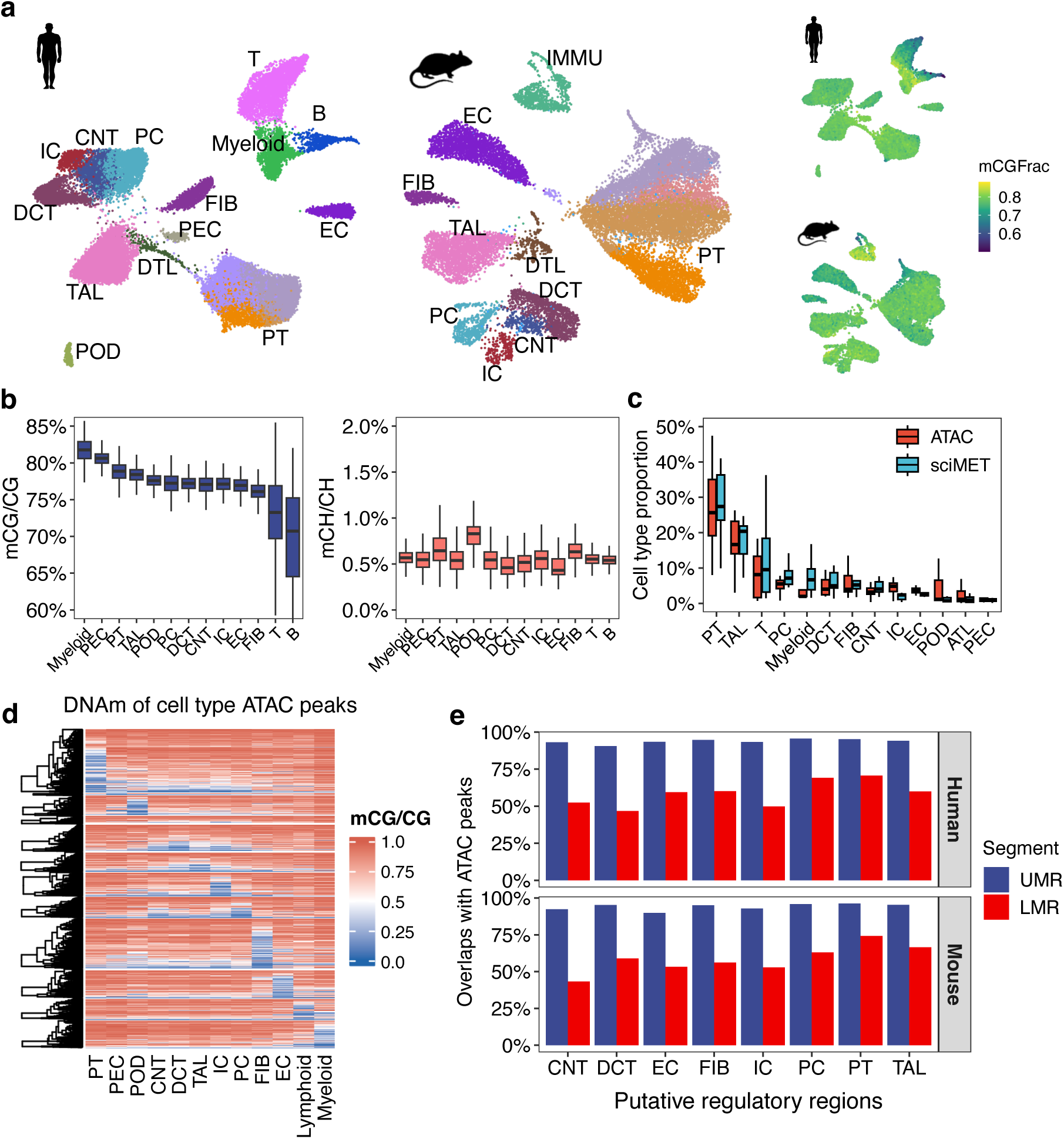
Single-cell DNA methylome analysis characterizes kidney cell type epigenomes and conservation across species. a, UMAP visualization of human and mouse kidney single-cell DNA methylomes. Clustering is based on genome-wide mCG profiles aggregated in non-overlapping 20-kb bins. **b,** Boxplots summarizing per-cell global methylation of mCG (upper) and mCH (bottom) for each kidney cell type. **c,** Cell-type composition of human kidney samples inferred from matched single-cell DNA-methylome and single-cell 10X Multiome (RNA + ATAC) datasets. **d,** Heatmap of mCG at cell-type-specific open-chromatin peaks, with peaks defined from matched human single nucleus 10x Multiome data. **e,** Fraction of ATAC-seq peaks that overlap putative regulatory elements: unmethylated regions (UMRs, promoter-like) and low-methylated regions (LMRs, enhancer-like).

We investigated the relationship between chromatin accessibility and DNA methylation at cell-type-specific regulatory elements using the matched single-cell multiome data. While, the cellular compositions from both modalities showed strong concordance (Spearman’s rho = 0.72, Fig. 1c), sciMET detected a higher proportion of immune cells, particularly myeloid lineages. This discrepancy may be attributed to the possibility that snRNA-seq under-detects immune populations, which can have lower nuclear RNA content. As expected, cell-type-specific ATAC peaks displayed clear hypomethylation in their corresponding cell type (Fig. 1d). To further characterize regulatory elements, we segmented the genome based on methylation patterns (Methods), identifying unmethylated regions (UMRs, <10% methylation, CpG rich) and low-methylated regions (LMRs, 10-50% methylation, CpG poor). These regions are known to be characteristic of promoters and enhancers, respectively ^20^. A majority of UMRs (92.6%) overlapped with cell-type-specific ATAC peaks, which is consistent with the nature of open and active promoters (Fig. 1e). In contrast, a smaller proportion of LMRs (54.8%) overlapped with ATAC peaks, suggesting greater variability in the relationship between methylation and accessibility within distal regulatory elements. Collectively, this multimodal integration highlights the concordance between chromatin accessibility and DNA methylation at promoters while capturing their distinct patterns at distal regulatory elements, establishing a rigorous high-resolution framework for characterizing kidney epigenomes.

### Epigenetic conservation and divergence in kidney cell types

We jointly analyzed human and mouse single-cell methylomes to quantify evolutionary conservation. To address sparse measurements, we aggregated each cell with its 50 nearest neighbors into meta-cells (Methods). These meta-cells were then profiled across 40,284 one-to-one orthologous 5 kbp bins, and the resulting datasets were integrated using Harmony ^21^. Following batch correction, human and mouse nuclei were co-clustered by cell type (Fig. 2a). Methylation patterns at canonical marker genes were consistent across homologous cell types. Leiden clustering in the integrated space identified 13 consensus kidney lineages. This was achieved using a reference panel of 2,180 conserved hypermethylated or hypomethylated regions shared between species (Fig. 2b and Supplementary Table 3). Overall, the cell type composition was largely similar between species, with abundance differences potentially due to differential sampling.

**Fig. 2.**
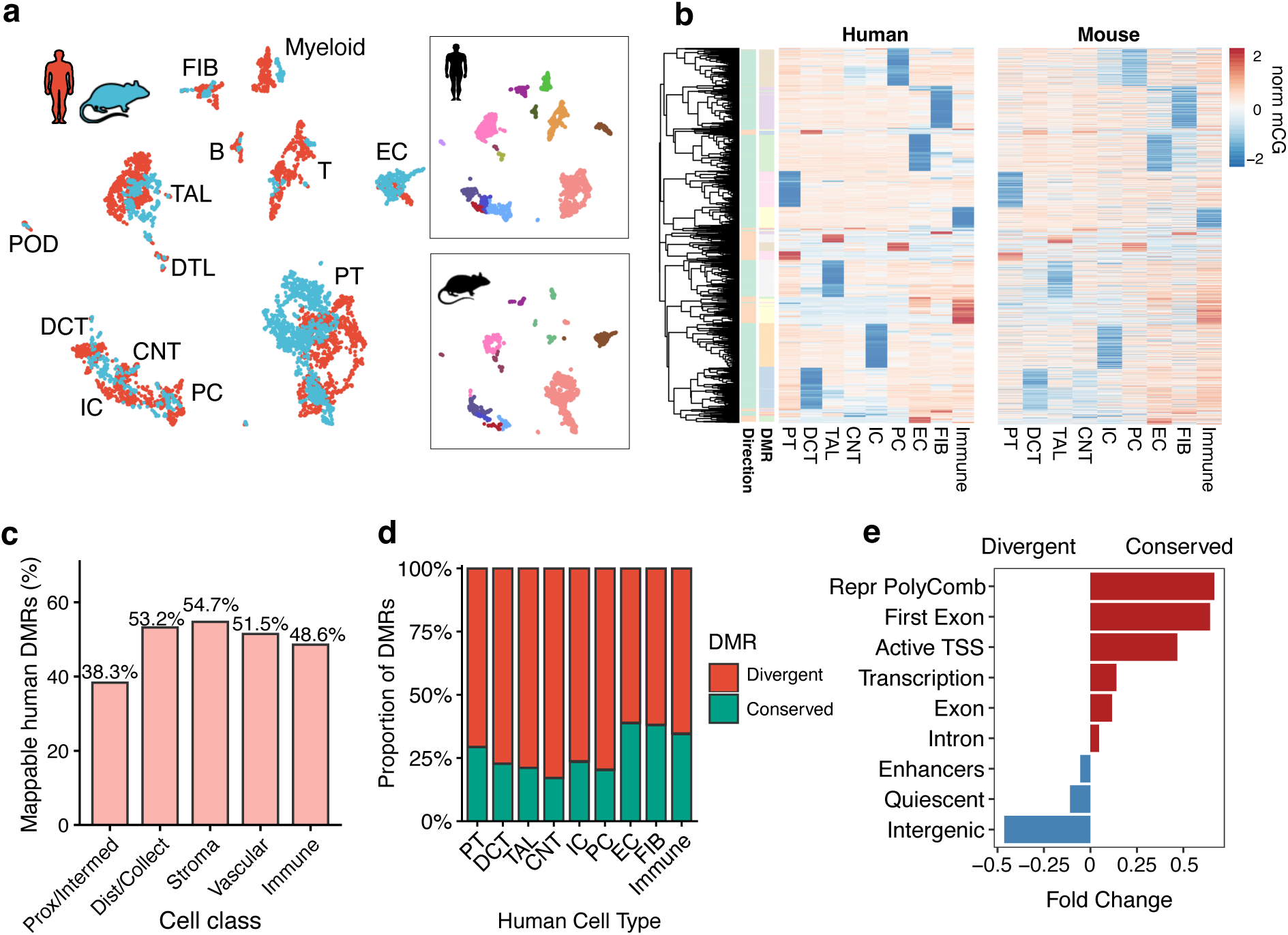
Comparative analysis reveals accelerated epigenetic evolution in kidney epithelium. a, UMAP visualization of the integrated human and mouse single-cell DNA methylation atlas, showing successful alignment of homologous cell types across species. **b,** Heatmaps of normalized methylation (z-score) at human cell-type DMRs (rows) and their orthologous mouse regions. **c,** Proportion of DMRs that was mappable between species. **d,** Fraction of human cell-type DMRs that were epigenetically conserved in mouse or species-specific (divergent). **e,** Differential enrichment of conserved versus divergent proximal tubule (PT) DMRs across functional annotations.

Analysis of differential methylation across human kidney cell types revealed 208,049 cell-type specific differentially methylated regions (DMRs). Whole methylome sequencing, although sparse at each cell, proved advantageous, as only 16.4% of these human DMRs overlapped with CpGs on the Illumina 850K array, suggesting that previous studies based on methylation arrays missed a larger fraction of regulatory elements in the genome. Of the cell-type DMRs in human, 138,110 regions were mappable to the mouse genome, with 57,259 aligning to one-to-one orthologous loci across species (Fig. 2c). We observed substantial conservation of cell-type-specific methylation states among these orthologous DMRs, with 34% of human DMRs retaining the same state in mice. However, epithelial cell types displayed significantly greater inter-species methylation divergence compared to non-epithelial populations (Fig. 2d).

This divergence was associated with lower underlying sequence conservation, which we observed particularly in DMRs from proximal and intermediate tubule epithelial cells (Fig. 2c). This observation aligns with previous cross-species studies showing greater species-specific divergence in epithelial populations compared to neural, muscle, and immune lineages, likely reflecting faster epigenetic evolution at distal regulatory elements in epithelia ^22^.

Analysis of differential methylation between species revealed distinct evolutionary patterns. Conserved cell type DMRs were enriched at transcription start sites and Polycomb repressed chromatin states, indicating active silencing of alternative lineage programs is an evolutionarily conserved mechanism for maintaining kidney cell identity across species. In contrast, species-specific DMRs were predominantly located in distal, lowly-methylated enhancer and intergenic regions (Fig. 2e).

In PT cells, both conserved and non-conserved DMRs showed enrichment for HNF4A and PPARA transcription factor binding motifs, highlighting their consistent roles in PT identity across mammals. However, AP-1 and MYC family transcription factor motifs were exclusively enriched in species-divergent PT DMRs, suggesting species-specific differences in the regulation of stress response, proliferation, and metabolic pathways (Supplementary Table 4). Functional annotation using GREAT ^23^ linked conserved PT DMRs to core nephron functions such as ‘organic anion transport,’ ‘urate transport,’ and ‘cellular amino acid metabolic processes’ (Supplementary Table 5). Conversely, the divergent PT DMRs were enriched for genome maintenance and stability processes closely associated with aging and cellular senescence, including ‘negative regulation of telomeric DNA binding,’ ‘negative regulation of histone H2A K63 linked ubiquitination,’ and ‘protein repair.’ They also showed enrichment for terms related to broader developmental patterning, including neural development, indicating weaker evolutionary constraints in the kidney or context-specific roles in other tissues (Supplementary Table 5).

Collectively, these findings indicate that evolutionarily stable differential methylation governs core renal physiology, while divergence in enhancer elements is enriched for chromatin maintenance and stress response pathways. These characteristics may contribute to the increased susceptibility of human epithelial cells to age-related epigenetic drift and disease-associated remodeling.

### Aging drives cell-type-specific epigenetic remodeling in the mouse kidney

Our analysis of the mouse kidney methylome captured a strong aging signature, as an epigenetic clock based on global DNA methylation accurately differentiated young (4 months) from aged (20 months) mice (Fig. 3a). We first characterized the methylation patterns of putative regulatory regions (UMRs and LMRs) across cell types. This revealed that regions with cell-type-specific hypomethylation were enriched in LMRs, while regions shared across cell types were enriched in UMRs (Fig. 3b,c). We then investigated whether aging selectively impacts these elements. Indeed, cell-type-specific UMR/LMRs exhibited significantly greater hypermethylation (Fig. 3d and Supplementary Fig. 3) and more pronounced age-related methylation shifts compared to their shared counterparts (Fig. 3e and Supplementary Fig. 4). This suggests that the regulatory elements maintaining specialized cell identities are intrinsically more susceptible to age-related decay than those governing core, shared cellular functions.

**Fig. 3.**
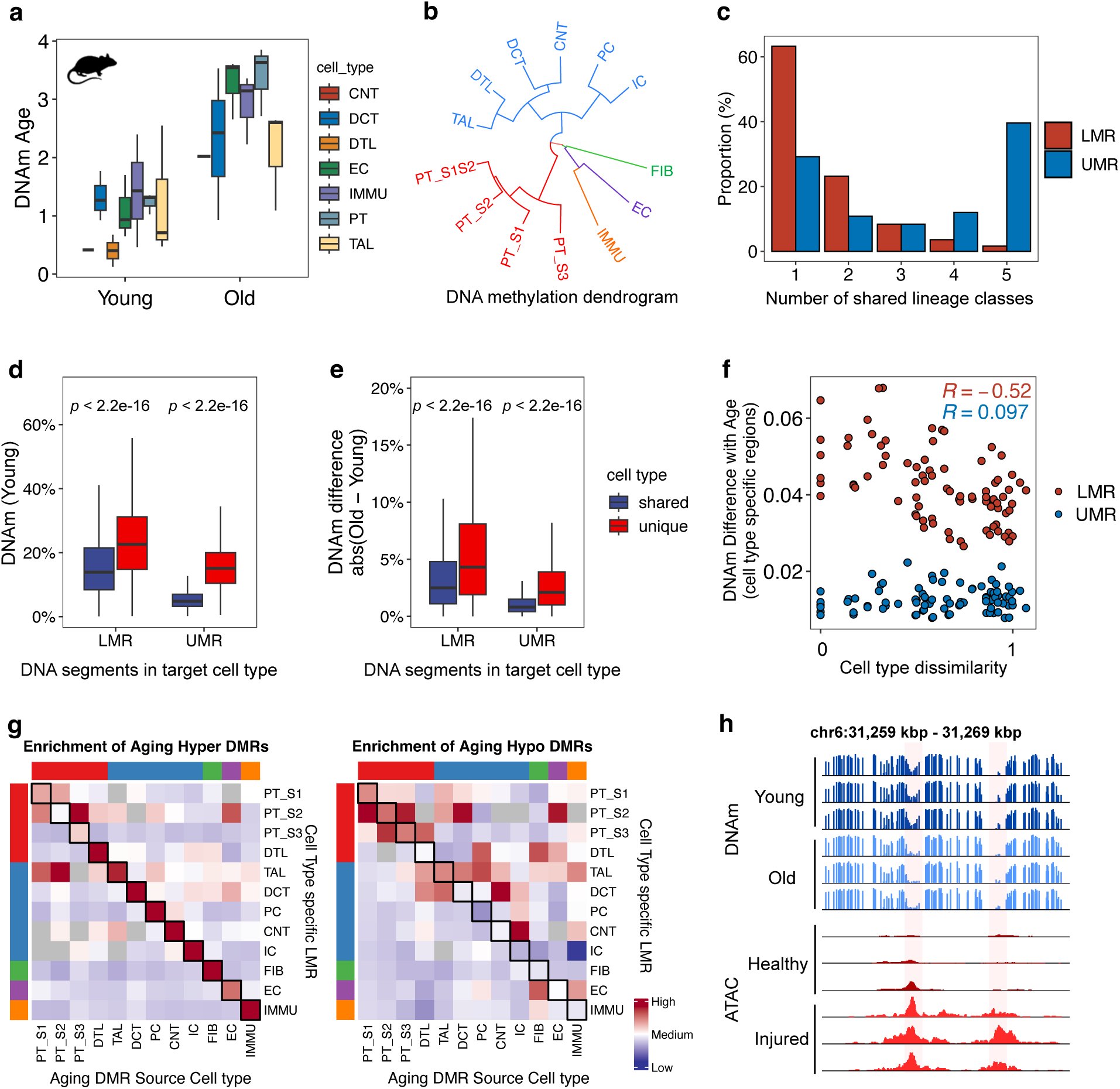
Age-related methylation changes selectively affect cell type-specific cis-regulatory regions, occurring disproportionately in regulatory regions specific to closely related cell types. a, Epigenetic-clock estimates for kidney cell-type pseudo-bulk methylomes from young (4 months) and aged (20 months) mice. **b,** Unsupervised hierarchical clustering of cell types based on genome-wide CpG methylation. Branch colors denote broad lineage classes. **c,** Distribution of hypomethylated elements across cell types. Bars show the proportion of low-methylated (LMR) or unmethylated (UMR) regions shared by lineage class colored in **b** (1 = lineage-unique; 5 = shared by all cell types). **d,** Box plots of mCG levels for LMRs and UMRs that are lineage-unique versus lineage-shared (data pooled across cell types; full breakdown by cell type in Supplementary Fig. 3). **e,** DNA methylation differences between age groups in lineage-specific versus lineage-shared LMRs and UMRs (a full breakdown by cell type in Supplementary Fig. 4). **f,** Correlation between inter-cell-type methylation distance and age-associated methylation differences. For a given target cell type, we first defined its cell-type-specific LMRs/UMRs. Then, for every other cell type, we calculated the average DNA methylation difference between age groups (old vs. young) at these specific loci. The cell-type methylation distance is quantified as the branch distance in the dendrogram shown in (b). Panels for each target cell type are shown in Supplementary Fig. 5. **g,** Heatmap of fold-enrichment for aging DMRs within cell-type-specific regulatory regions. Enrichment is calculated relative to 100 CpG- and GC-matched random sets; red denotes enrichment, blue denotes depletion. **h,** Genome browser snapshot of a 10 kbp window showing aging-associated hypomethylation alongside chromatin accessibility tracks for proximal tubules in healthy and injured states from a mouse ischemia-reperfusion injury (IRI) model.

To understand how age-related methylation changes correlate with cell-type relatedness, we quantified cell-type-specific UMR and LMR methylation shifts in their unassociated cell types (Methods). A strong inverse relationship was observed between inter-cell-type methylation distance and the magnitude of aging-associated methylation change, indicating that regulatory regions defining closely related cell types were more vulnerable to age-associated epigenomic dysregulation (Fig. 3f and Supplementary Fig. 5). This pattern was exclusive to LMRs, suggesting that the epigenetic signatures of distal enhancers were more susceptible to age-related changes than those of more broadly active promoters. Additionally, regions that became dysregulated with age tended to be closer to regulatory elements, implying that the chromatin context may facilitate methylation remodeling at already accessible sites (Supplementary Fig. 6).

These results suggest that regulatory elements potentially underlying distinct phenotypic functionalities of closely related cell types might be more susceptible to age associated dysfunction.

Differential methylation analysis revealed that age-associated DMRs were significantly enriched in cell-type-specific cis-regulatory regions (Fig. 3g), suggesting these loci are preferential targets of age-related epigenetic drift. This enrichment reflects a progressive loss of epigenetic identity. Specifically, cell types gained methylation at regions that were hypomethylated in young mice and concurrently showed age-associated hypomethylation at cis-regulatory regions specific to other cell types. This divergence of epigenetic cell type identities was further evident at the PT subtype level, where hypomethylated DMRs in one segment were enriched in the LMRs of an adjacent PT segment (Fig. 3g).

Our findings indicate that the regulatory elements defining cell identity are uniquely susceptible to aging-associated dysregulation. To directly link the aging epigenetic signatures to kidney pathology, we analyzed single-nucleus ATAC-seq data from a mouse model of renal ischemia-reperfusion injury (IRI) to identify differentially accessible regions that define the adaptive and maladaptive PT repair states ^24^.

Remarkably, we found that DNA methylation changes from normal aging were significantly enriched within these injury-induced chromatin accessible peaks (fold enrichment = 2.95, *P* < 0.01, permutation test; Fig. 3h). This overlap demonstrates that genomic loci susceptible to age-related epigenetic drift are also hotspots for regulatory reprogramming during acute injury.

### Kidney tubular epithelial cells are the primary drivers of accelerated epigenetic aging in disease

To investigate whether CKD is linked to accelerated epigenetic aging, we compared single-cell methylomes from CKD patients with those from age-matched healthy controls. Our initial analysis of sample-level pseudo-bulk methylomes revealed significant epigenetic age acceleration in CKD patients (*P* = 0.01, Wilcoxon rank-sum test). We further ensured the robustness of this finding by checking for technical bias through down-sampling reads and employing a principal component-trained version of the Horvath clock ^25^, which performs robustly with low-coverage methylome data. Age predictions remained accurate at shallow coverage (down to approximately 3X), supporting the reliability of the epigenetic age calculation (Supplementary Fig. 7).

Cell-type level analysis demonstrated that epigenetic age acceleration in CKD was dominated by epithelial cells (Fig. 4a). Specifically, epithelial cells from CKD patients showed an average epigenetic age acceleration of 4.8 years when compared to those from healthy controls (*P* = 2.3×10^−4^). In contrast, non-epithelial populations, including fibroblasts, endothelial cells, and immune cells, did not show significant acceleration in the same comparison (Fig. 4a). This epithelial-specific age acceleration appears to be due to cell-intrinsic epigenetic changes, rather than shifts in cell type composition, as disease samples showed similar epithelial proportions while showing increases in fibroblasts and immune cells (Fig. 4b).

**Fig. 4.**
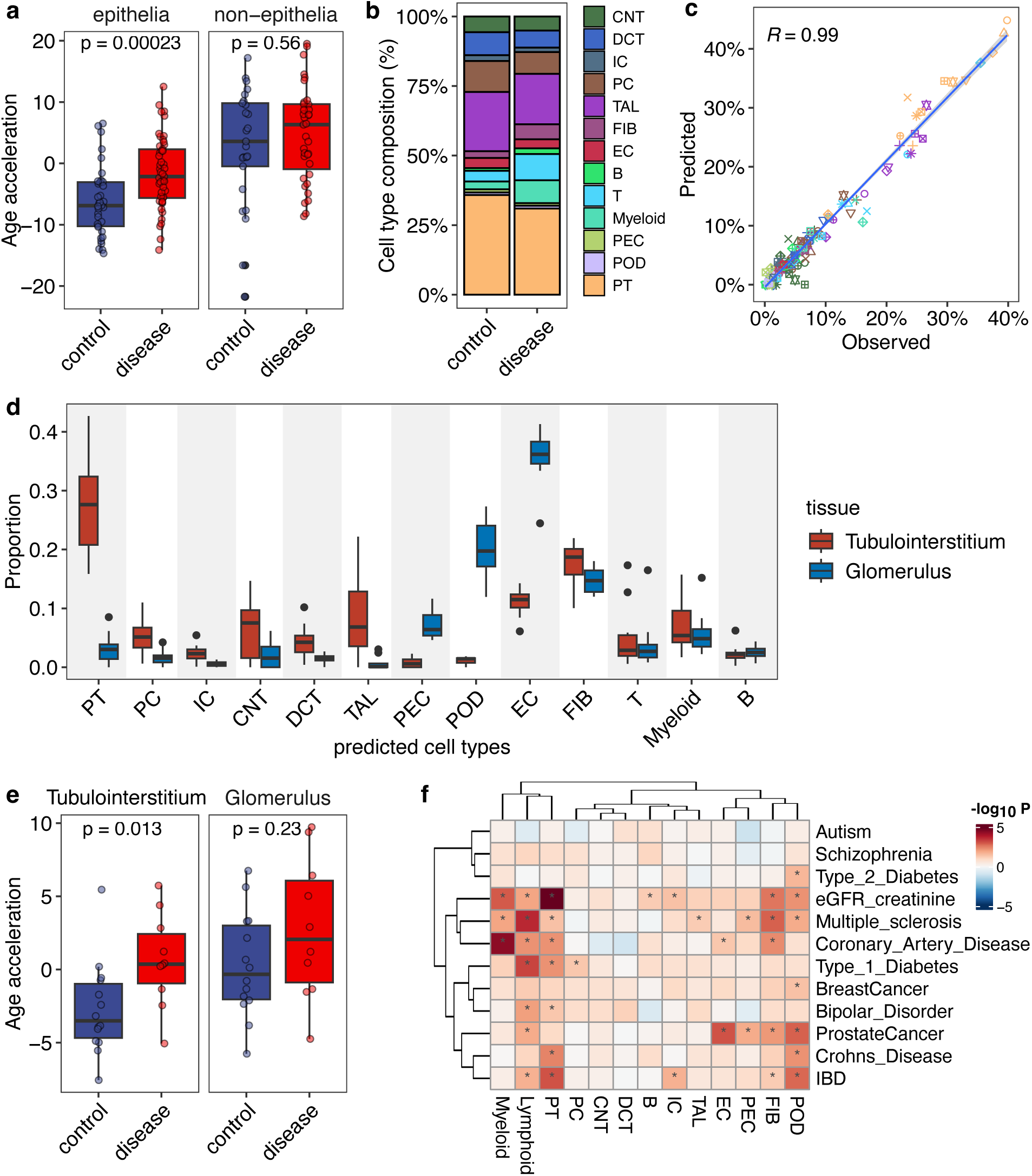
Accelerated epigenetic aging in tubular epithelial cells is a hallmark of chronic kidney disease (CKD). a, Cell-type-specific epigenetic age acceleration in CKD patients versus healthy controls. Age acceleration, defined as the residual difference between predicted DNA methylation age and chronological age, was calculated from sample-level cell type pseudo-bulk methylomes. The analysis highlights that accelerated aging is most prominent in epithelial cell types. A principal-component (PC)-based implementation of Horvath multi-tissue clock was used for age prediction. **b,** Stacked bar charts comparing cell-type composition (% nuclei) between control and CKD kidneys, illustrating expansion of fibroblast and immune cells in disease. **c,** Accuracy of the methylation-based deconvolution model. The plot shows the correlation between the predicted cell-type proportions and the observed proportions derived from single-cell annotations, using a leave-one-out cross-validation approach. **d,** Application of the validated deconvolution algorithm to an independent bulk kidney tissue cohort. The analysis estimates cell-type composition in whole-genome bisulfite sequencing (WGBS) data from laser-capture microdissected glomerular (GLOM, n=15) and tubulointerstitial (TI, n=15) compartments. **e,** Comparison of age acceleration in bulk GLOM and TI tissues from control and CKD patients. Consistent with single-cell findings, the TI compartment exhibits significant age acceleration in CKD. **f,** Partitioned heritability analysis showing the enrichment of genetic risk for complex traits within cell-type-specific DMRs (extended by +/- 25kb). The plot displays the statistical significance of enrichment. Significant associations (FDR < 0.05) are marked with an asterisk. DMRs specific to Proximal Tubule (PT) cells are highly enriched for heritability of the kidney function trait eGFR.

To validate our findings, we used bulk whole-genome bisulfite sequencing (WGBS) data generated independently from laser-microdissected kidney tubulointerstitial (TI, n=15) and glomerular (GLOM, n=15) tissues ^26^. We performed cell type deconvolution using our single-cell methylome atlas as a reference (Pearson’s r=0.97; Fig. 4c) to examine cell type proportions within the bulk methylome data. This deconvolution approach identified an expected enrichment of tubular epithelial cells in TI samples and of podocytes, endothelial cells, and parietal epithelial cells in glomerular samples (Fig. 4d). Epigenetic clock estimates from these deconvoluted cell populations confirmed that age acceleration in CKD is most pronounced in tissues enriched for tubular epithelial cells (Fig. 4e).

Building upon our previous finding that altered PT states of unresolving or maladaptive repair are associated with chronic disease progression ^9,10^, we sought to link the regulatory elements of kidney cell types to genetic kidney disease risk. Using stratified linkage disequilibrium score regression (S-LDSC) ^27^, we observed a strong enrichment of PT-specific DMRs for the genetic heritability of estimated glomerular filtration rate (eGFR) (Fig. 4f). Notably, we also observed enrichments for risk variants associated with Type 1 Diabetes and Inflammatory Bowel Disease (IBD). Both are systemic conditions with established links to kidney pathology ^28–30^. This enrichment was also evident in DMRs that define the transition from a healthy to an altered state (FDR = 0.004), thereby directly linking epigenetic dysregulation of this cell state to genetic risk variants for CKD.

### Altered proximal tubule cells exhibit epigenetic instability and recapitulate transcriptomic signatures of aging

To define the epigenomic changes of this pathological state, we compared altered- and healthy-state PT cells from the same CKD donors, an approach that accounts for inter-individual genetic variation (Methods). This revealed extensive epigenetic alterations between the cell states (Fig. 5a), with a trend towards gain of methylation, as 63% of the differential methylation was hypermethylated (Supplementary Table 6). Functional annotation identified the significant enrichment of terms including “response to laminar fluid shear stress” (FDR = 2 x 10^-11^) and “regulation of histone deacetylation,” (FDR = 7.8 x 10^-9^) implying a significant shift of epigenetic landscape between the cell states (Supplementary Table 7). Furthermore, these methylation changes were significantly enriched within the PT-specific DMRs that defined healthy tubular identity (fold-enrichment = 1.84, *P* < 0.01), indicating that regulatory regions of steady-state PT were specifically dysregulated in disease.

**Fig. 5.**
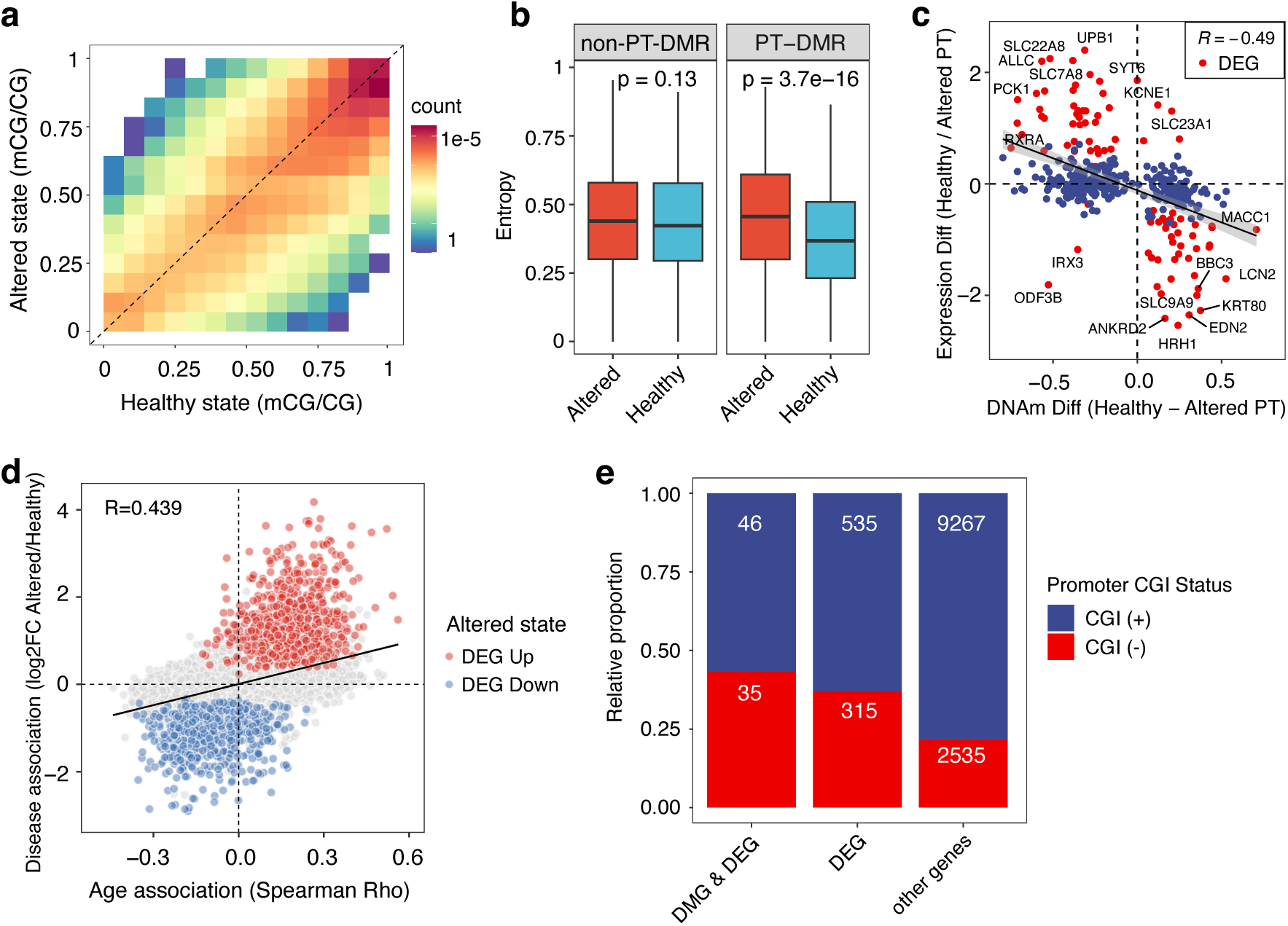
Altered-state proximal tubule cells show a loss of locally coordinated methylation, a disruption that is significantly enriched at genes lacking CpG islands. a, Density plot comparing per-CpG methylation levels between healthy and altered PT cells. The average DNA methylation difference at each CpG locus was computed within the same donor. **b,** Quantification of intra-population methylation heterogeneity (epiallelic entropy). Within the altered PT cell population, PT-specific DMRs exhibited significantly higher methylation entropy than DMRs specific to other cell types, indicating greater stochastic variability at cell type specific regulatory regions. **c,** Scatter plot correlating promoter DNA-methylation change with log2 fold-change in gene expression between healthy and altered PT states. Genes that were significant (FDR < 0.05) both as differentially methylated (DMG) and differentially expressed (DEG) are highlighted in red. **d,** Scatter plot showing the relationship between age-association (Spearman’s rho, X-axis) and differential expression in the altered PT state (log2 fold change, Y-axis). Positive Y-values denote higher expression in the altered state PT. Differentially expressed genes (FDR < 0.05) are colored by direction (Red: Up in Altered; Blue: Down in Altered). **e,** Barplot displaying the proportion of genes containing or lacking CpG islands, stratified into three groups: genes that were both DMG and DEG, genes that were DEG only, and all other genes.

Beyond average methylation changes, altered-state PT cells exhibited a profound increase in epigenetic noise. Single-chromosome co-methylation analysis, which measures the consistency of methylation states across neighboring CpG sites on the same DNA molecules ^31^, revealed a significant reduction in methylation concordance in altered-state cells, accompanied by increased local methylation entropy (Fig 5b). This loss of local epigenetic order, an indication of loosen local epigenetic control, was most pronounced at PT-specific regulatory regions, mirroring our findings in the aging mouse kidney.

Next, we investigated whether the cell state change disrupts the activity of key transcription factors (TFs) that maintain PT identity. Using the matched single-cell ATAC-seq data, we identified TF binding motifs with enriched accessibility in healthy PT cells, including those for the master regulators of the HNF, RXR, and PPAR families (Supplementary Fig. 8a). In altered-state PT cells, these lineage-defining TF motifs showed significant gain of methylation (Supplementary Fig. 8b), suggesting that the epigenetic silencing of this master regulatory network might be a key mechanism driving the loss of PT cell identity in CKD.

These epigenetic alterations had direct functional consequences on local gene expression. We observed a significant negative correlation between methylation at DMRs and the expression of neighboring genes (Spearman’s rho=−0.49; Fig. 5c). To further investigate if the pathological state reflects an acceleration of normal biological aging, we analyzed age-associated transcriptional changes using single-nucleus transcriptome data from 72 healthy kidneys (ranging from 20 to 90 years of age). We found that age-associated dysregulation was highly cell-type specific, with proximal tubules exhibiting a distinct bias toward gene upregulation over downregulation (Supplementary Fig. 9). The top age-upregulated genes in healthy PT cells included pro-inflammatory chemokines (*CXCL1*), senescence-associated factors (*IGFBP6*), and ion channels (*KCNB2*) (Supplementary Fig. 10). Notably, the gene expression changes associated with normal aging showed a significant positive correlation with the transcriptional shifts defining the altered PT cell state (Fig. 5d), indicating that the pathological cell state in CKD recapitulates an accelerated aging phenotype at the molecular level.

Finally, we found that genes differentially methylated and differentially expressed in the altered state were enriched for those lacking CpG islands (CGI-less) (Fig. 5e). CGI-less genes are preferentially located within repressive heterochromatin, particularly lamina-associated domains, where their misexpression has been identified as a driver of aging-associated degeneration ^32,33^. Consequently, the dysregulation of CGI-less genes in altered-state cells implies a breakdown of repressive heterochromatin structures and suggests a potential disruption of higher-order chromatin architecture.

### Differential methylation in altered state is associated with the upregulation of genes that define the disease-associated niche

To understand how methylation-associated gene expression changes relate to human disease pathology, we mapped the spatial transcriptomics profiles of 5000 genes with the 10X Xenium platform on either healthy, acute kidney injury (AKI) or CKD tissues (Supplementary Figs. 11,12). Integration with the Human Kidney Atlas (HKA) ^9,10^ resolved cell types spanning cortical to medullary structures, as well as adaptive or maladaptive PT states associated with resolving or unresolved repair (Supplementary Fig. 11). These cell types were further organized into niches based upon their spatial colocalizations (Fig. 6a, Supplementary Fig. 12a), to resolve normal renal structures (e.g. Niche 10 renal corpuscles) as well as distinct early (Niche 8) or failed (Niche 6) epithelial repair niches. Consistently these niches showed a switch from healthy (HNF4A) to adaptive (NFkB, SOX4) PT transcription factor activities (Supplementary Fig. 12b). Furthermore, the early PT repair Niche 8, enriched in the AKI sample showed increased early stress response TF activities (ATF4, JUN, BACH2) and increased activity of the E2F pathway (Fig. 6b,c, Supplementary Fig. 12b). Alternatively, the failed repair niche, enriched in CKD, showed higher TF activities associated with the failed repair PT state (BHLHE40, MYC, MITF) ^10^ and elevated Wnt signaling that has been associated with fibrosis (Fig. 6b-c, Supplementary Fig. 12b).

**Fig. 6.**
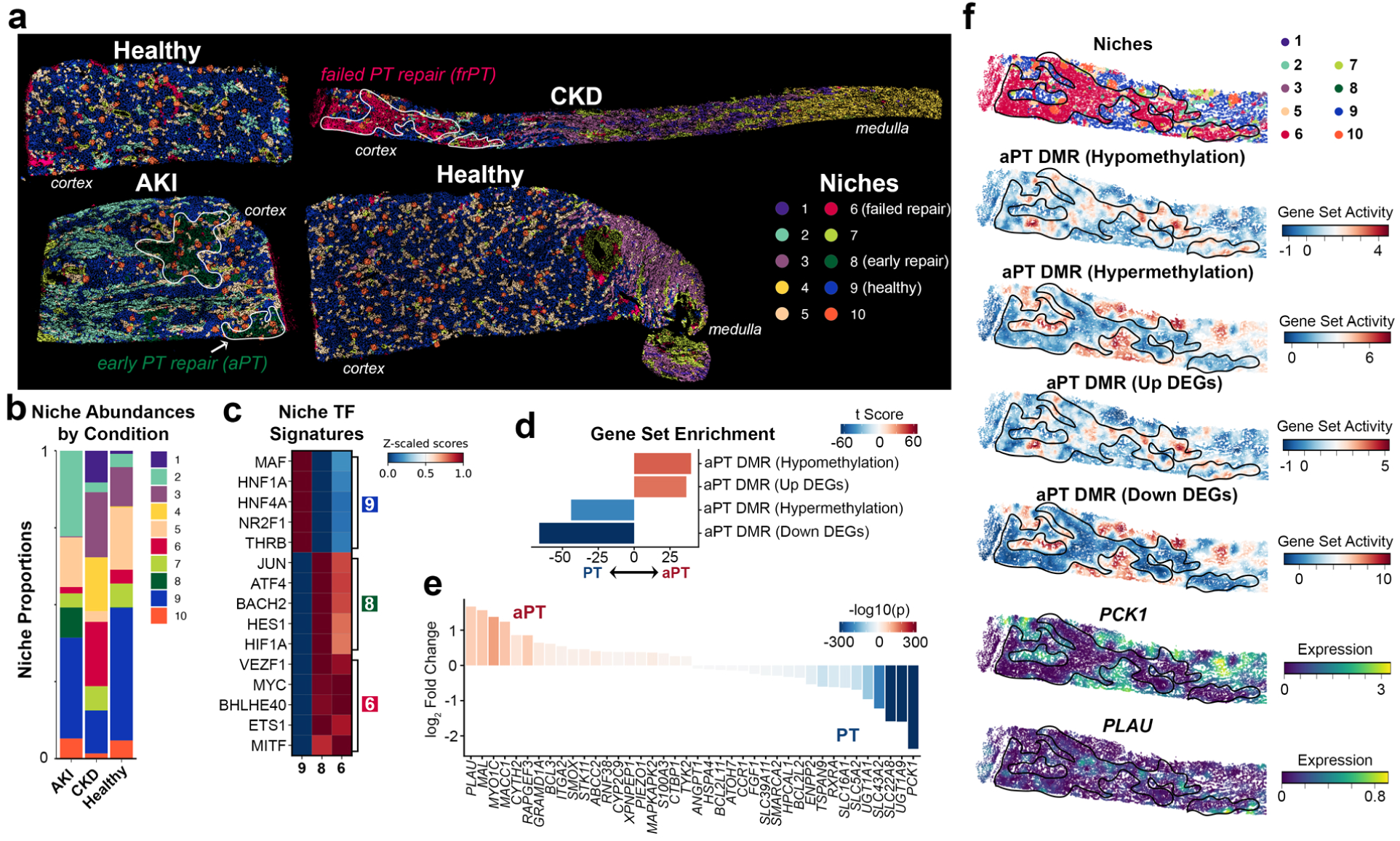
Spatial localization of altered state PT DMR-associated genes to pathological niches. a, Xenium spatial niches based on cell type or state co-localizations mapped to healthy reference tissues (HRT), or tissues associated with Acute Kidney Injury (AKI) or chronic kidney disease (CKD). Areas associated with early PT repair states (AKI) or late failed repair states (CKD) are indicated. **b,** Barplot showing proportion of each niche for HRT, AKI and CKD samples shown in (a). **c,** Heatmap of transcription factor (TF) associated target gene expression across niches associated with healthy (9), early repairing (8) or failed repair (6) PT cells. TF-gene weights were derived from previously defined gene regulatory networks associated with PT cell state shift trajectories (Methods). **d,** Waterfall plots showing t-scores comparing associated gene set enrichment scores between cells from each group (healthy PT vs. altered PT) using Xenium data. **e,** Waterfall plots showing t-scores comparing expression of DMR-associated genes from (d) between each group (healthy PT vs. altered PT). f. Spatial mapping of gene set scores from (d) and gene expression values for the top two differentially expressed genes (e) to the CKD sample shown in (a). Areas associated with the failed repair PT niche (6) shown in the top panel are highlighted.

Gene set enrichment analysis (GSEA) confirmed an enrichment of altered-state PT hypomethylated genes in Xenium altered PT cells, with altered-state PT hypermethylated genes enriched in the healthy state (Fig. 6d). This is consistent with trends for genes also found to be differentially expressed between these states in the 10X Multiome data (Fig. 6d). PCK1, encoding a key component of the gluconeogenesis metabolic pathway found to be critical for normal PT function, was found as a top gene that was hypermethylated and downregulated in altered-state PT cells (Fig. 6e) ^10^.

Alternatively, genes associated with cell polarity and adhesion were hypomethylated and upregulated in altered-state PT cells (PLAU, MYO1C, CYTH2), consistent with the mesenchymal-like shift associated with this state ^10^. Spatial mapping of DMR gene signatures further confirmed their differential enrichment between healthy (Niche 9) and pathology-associated (early repair Niche 8 and failed repair Niche 6) niches (Fig. 6f, Supplementary Fig. 12c).

### PT-specific chromatin compartments are vulnerable to reorganization in the altered state

To determine if the observed local epigenetic instability and dysregulation of CGI-less genes are linked to a global reorganization of higher-order chromatin structure, we performed scMethyl-Hi-C ^17^, which simultaneously captures both 3D chromosome conformation and the DNA methylome within the same cell. We generated 3,072 high-quality single-cell profiles on a healthy reference donor, yielding over 300 million long-range chromosomal contacts, allowing us to define the baseline 3D chromatin compartments characteristic of kidney cell types. Using the DNA methylome component of this assay, we accurately annotated the cell type of each nucleus by integrating the data with our primary sciMET reference atlas, observing no significant batch effects between the two modalities (Fig. 7a). Furthermore, we demonstrated that 3D chromatin contact information alone was sufficient to separate major kidney cell types, highlighting that distinct genome folding patterns are a defining feature of cell identity (Fig. 7b).

**Fig. 7.**
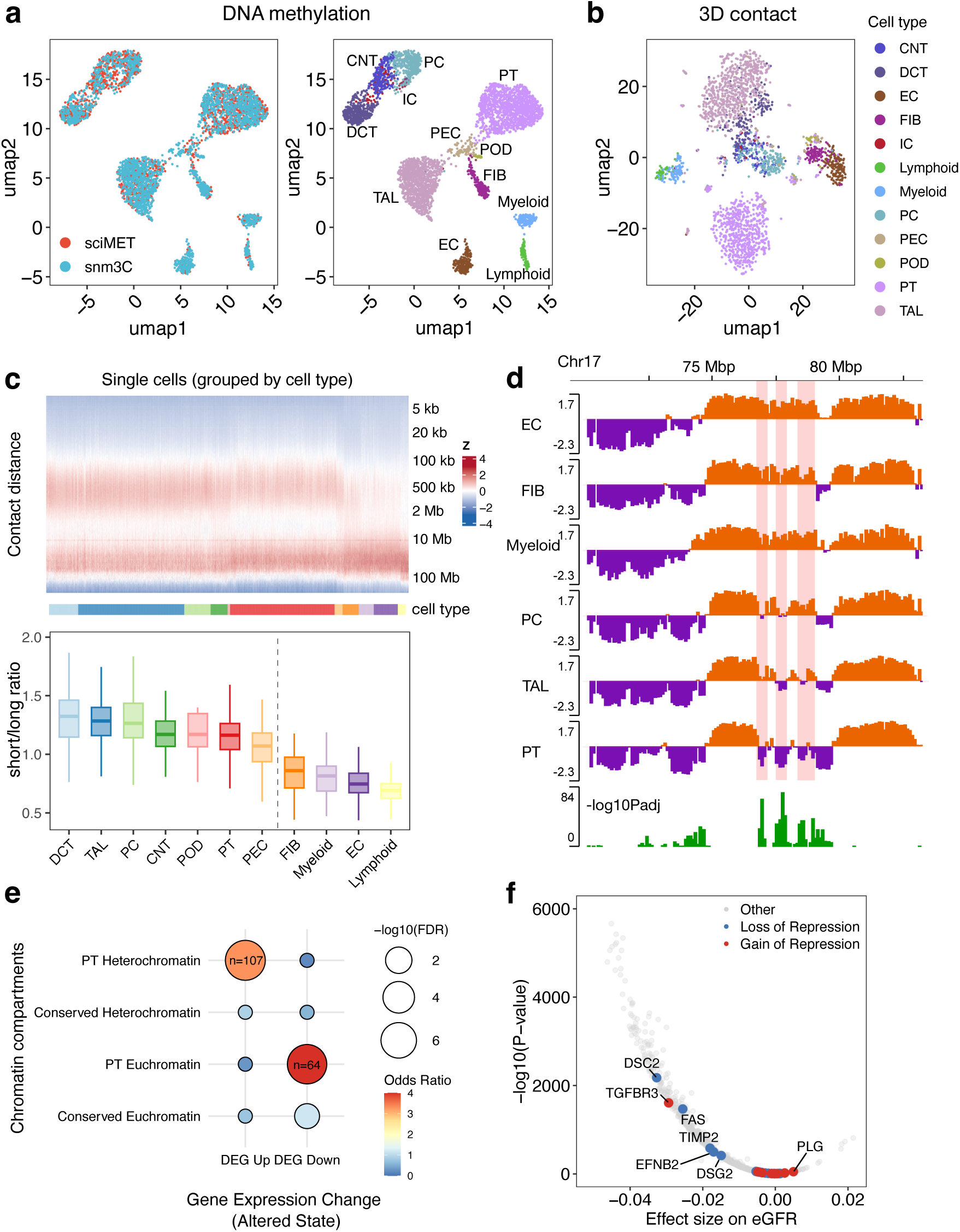
Single-cell methyl-Hi-C reveals that PT-specific chromatin compartments are vulnerable to reorganization in the altered cell state. a, UMAP visualizations demonstrating integration of the sciMET and scMethyl-Hi-C datasets. Cells from both assays co-clustered by cell type, indicating minimal batch bias after Harmony correction. **b,** UMAP visualization of single-cell Hi-C contact profiles, generated using 100kb genomic bins. Each point represents a single nucleus, colored by its annotated cell type in (a). **c,** Per-nucleus contact-distance profiles for scMethyl-Hi-C data. Heatmap rows represent normalized frequency of genomic distance between pairs; columns are single nuclei. Darker colors indicate higher contact frequency at a given distance. **d,** Example genome browser view of loci undergoing a chromatin compartment switch in PT cells (highlighted in red). The first principal component (PC1) track derived from Hi-C data was used to estimate putative euchromatin and heterochromatin compartments. **e,** Dot plot illustrating the enrichment of altered state differentially expressed genes (DEGs) within PT-specific versus conserved chromatin compartments. Dot size corresponds to statistical significance (-log10 FDR), and color intensity represents the magnitude of enrichment (Odds Ratio, Fisher’s Exact Test). **f,** Scatter plot of 1,463 plasma proteins from the UK Biobank Olink panel, showing their association with estimated Glomerular Filtration Rate (eGFR). The X-axis represents the effect size (Beta coefficient) of protein levels on eGFR, and the Y-axis represents statistical significance (-log10 P-value). Proteins were stratified based on the epigenetic stability of their encoding genes in PT cells.

Analysis of global chromatin organization revealed a fundamental structural distinction between epithelial and non-epithelial kidney cell types. Specifically, epithelial cells displayed a significantly higher ratio of short-range (up to 2 Mbp) to long-range (2-10 Mbp) contacts when compared to non-epithelial cell types (Fig. 7c). A high prevalence of short-range contacts is a hallmark of a well-organized genome, reflecting the dense packing of genes into discrete topologically associating domains (TADs) that facilitate precise local regulation ^34,35^. This suggests that epithelial cell identity relies more heavily on complex, precise local 3D folding, potentially making it more susceptible to structural disorganization compared to other lineages.

To determine if the 3D chromatin architecture of proximal tubule cells is disrupted during the transition to the altered state, we constructed cell-type-specific maps of euchromatin (active) and heterochromatin (repressed) compartments at 250 kb resolution. PT compartment scores were compared against the other kidney cell types (Methods), where PT-specific compartments were defined as chromatin state regions that were unique to the PT lineage (Fig. 7d). We found that genes upregulated in the altered state were significantly enriched within PT-specific heterochromatin compartments (Fig. 7e). This selective enrichment implies that the activation of disease-associated genes involves a specific breakdown of silencing at loci defining PT identity. Consequently, we annotated 107 PT-specific heterochromatic compartments as undergoing a loss-of-repression, and 64 PT-specific euchromatic compartments as undergoing a gain-of-repression (Supplementary Table 8). Importantly, the majority of these regions also exhibited expected concurrent DNA methylation changes (70% hypomethylation in loss-of-repression loci; 78% hypermethylation in gain-of-repression loci), validating our structural annotations.

Analysis of genes located within the loss-of-repression domains revealed multiple loci known to be involved in pathological tissue remodeling (Supplementary Table 8). The loss-of-repression domains included genes known to promote inflammation and fibrosis that are typically silenced in healthy epithelia. Notable examples include the senescence-associated secretory phenotype (SASP) chemokines (*CXCL1*, *CXCL2*, *CXCL3*, *CXCL6*) and profibrotic ECM components and receptors (*ITGA2*, *ANTXR1*).

Furthermore, we observed the de-repression of transcription factors linked to epithelial-mesenchymal transition (EMT), such as *HMGA2*, *ETS1*, and *BACH2*. This suggests that the erosion of the cell type repressive compartments activate transcriptional programs that drives the pathological dedifferentiation of the proximal tubule.

Finally, we investigated whether the de-repression of these genes has systemic clinical relevance. We cross-referenced our loss-of-repression gene list with the Olink plasma proteomics data from the UK Biobank ^36^, which provides paired measurements of circulating protein abundance and kidney function (eGFR) from 54,219 participants. We identified that secreted proteins encoded by the epithelial derepressed genes, specifically *DSC2*, *DSG2*, and *TIMP2*, show significant negative associations with eGFR in the general population (Fig. 7f). Collectively, these findings demonstrate that the disease-associated cell state changes are driven by the selective collapse of higher-order chromatin structures, activating pathogenic pathways that contribute to tissue fibrosis and systemic renal decline.

## Discussion

DNA methylation provides a stable record of cellular identity and stress, making it a critical layer for understanding chronic diseases. While accelerated epigenetic aging is a known hallmark of chronic disease ^37–41^, these analyses have largely been performed on bulk tissues, obscuring the contributions of individual cell populations and leaving the cell-type- and state-specific epigenetic programs that drive disease largely uncharacterized. Our study addresses this gap by using a single-cell DNA methylome to demonstrate that the accelerated epigenetic aging in CKD is driven almost exclusively by the tubular epithelium. We also found that eGFR-associated genetic risk variants are enriched in differential methylated regions, which is linked to cell state changes in the proximal tubule.

Analysis of single-nucleus transcriptomes from a healthy cohort revealed that the transcriptional changes defining the disease-associated altered cell state are highly correlated with changes occurring during normal aging. This strong correlation suggests that the pathological state represents an acceleration of the natural erosion of epigenetic cell type identity. We found that the loss of epigenetic fidelity disproportionately affects genes lacking CpG islands, which typically encode highly cell-type or tissue-specific genes ^33^. Because these lineage-specific genes are typically located in lamina-associated domains to maintain silencing, the age-associated erosion of these repressive structures makes them highly prone to abnormal activation ^32^.

Furthermore, we show that altered cell states in the proximal tubule are characterized by a multi-scale collapse of epigenetic identity, ranging from a loss of local methylation coordination to a disorganization of higher-order 3D chromatin architecture. Our scMethyl-Hi-C analysis reveals that this structural disruption is not random, but rather selectively targets PT-specific chromatin compartments. We observed a specific loss-of-repression at these lineage-defining structural domains. This disruption leads to the aberrant activation of pathogenic pathways, including SASP chemokines and EMT factors, that are normally silenced in healthy epithelia. The clinical relevance of this structural failure is underscored by our finding that secreted proteins encoded by these de-repressed genes (e.g. DSC2, DSG2, and TIMP2) are significant predictors of eGFR decline in the general population ^36^.

This high-resolution, cross-species kidney epigenome atlas represent a key resource to the community. Our comparative analysis of human and mouse methylomes revealed that while core renal cell identities are evolutionarily conserved, epithelial lineages, particularly the proximal tubule, exhibit significant epigenetic divergence. This divergence was most pronounced in distal regulatory elements linked to pathways such as stress response and genome maintenance. This suggests that although the basic metabolic functions of epithelial cells are conserved, the regulatory circuits governing their response to stress represent more recently evolved adaptations. This inherent regulatory divergence in epithelia may make them particularly susceptible to the epigenetic dysregulation associated with chronic disease.

Our study has several limitations. Firstly, the mouse models of aging and acute injury used in this research may not fully capture the intricate, multifaceted nature of human chronic disease. Future longitudinal studies are necessary to definitively establish a causal link between epigenetic aging and CKD progression. Secondly, our cross-species comparisons primarily focused on orthologous bins and mapped differentially methylated regions. Consequently, species-specific sequences and human genetic variations might introduce additional effects that were not entirely accounted for in this study. Lastly, due to the relatively sparse resolution of scMethyl-Hi-C, we likely underestimated the full extent of chromatin loop disruption. Despite these limitations, our study collectively provides valuable insights into the cell-type-level epigenomes in the kidney, providing a foundational resource for future investigations and deciphering genome wide methylation-based physiological decline of kidney function.

## Methods

### Sample collection

Human kidney samples were obtained from adult individuals, including healthy controls and CKD patients (nephrectomies or tumor-adjacent normal tissue), ages ranging from 30-75 years old (Supplementary Table 1). Mouse kidney samples were collected from young (4 month) and old adult (20 month) C57BL/6 mice (Supplementary Table 2). All samples were processed into single nuclei suspensions by tissue mincing, enzymatic digestion, and nuclear isolation. We isolated nuclei to ensure efficient DNA recovery and to facilitate combinatorial indexing library prep. All procedures were approved by relevant institutional review boards and animal care committees. Human kidney samples were collected by the Kidney Translational Research Center (KTRC) as part of the Human Biomolecular Atlas Program (HuBMAP). The study protocol was approved by the Washington University Institutional Review Board (IRB #201102312). Informed consent was obtained from all participants, including living patients undergoing partial or total nephrectomy and deceased donors whose organs were declined for transplantation.

### Single-cell methylome library preparation

Nuclei were isolated from frozen OCT-embedded tissue sections. For human samples, 16-23 sections (10 µm thickness) were collected per donor, while for mouse samples, 5-7 sections were pooled per animal. Following isolation, we performed single-cell DNA methylation sequencing using the Scale Biosciences sciMET protocol ^18^. In brief, nuclei were first distributed into a 96-well plate and subjected to in situ Tn5 transposase tagmentation with indexed adapters (each well with a unique Tn5 index). The transposase fragmentation integrates adapter sequences into the genomic DNA of each nucleus. Next, nuclei from all wells were pooled, thoroughly mixed, and then redistributed by FANS into 12 96-well plates. In the redistributed plate, we carried out bisulfite conversion followed by indexed primer extension. This two-level indexing design uniquely tags DNA from each individual nucleus with a combinatorial barcode, allowing sequencing of many single-cell methylomes in one batch. Libraries were PCR-amplified with standard Illumina primers and purified.

### Sequencing and data analysis

sciMETv2 and single-cell methyl-Hi-C libraries were sequenced on an Illumina NovaSeq S4 platform using 150 bp and 100 bp paired-end reads, respectively. Reads were aligned to the human (hg38; RefSeq Feb 3, 2022) or mouse (mm39; RefSeq Jan 24, 2020) reference genomes using Bismark ^42^ aligned with Bowtie2 ^43^. Following deduplication, we obtained a mean coverage of approximately 0.8 million unique CpG sites per cell. Cells with <200,000 covered CpG sites were filtered out for downstream analysis. Fractional methylation levels were calculated at each CpG site for every cell.

Dimensionality reduction was performed using principal component analysis (PCA) on highly variable methylated regions, followed by Uniform Manifold Approximation and Projection (UMAP) ^44^ for visualization.

Cell type annotation was performed by assessing differential methylation patterns at established cell-specific marker loci (Supplementary Fig. 1). To achieve high-resolution annotation of proximal and distal tubule subsets, we integrated our methylation data with a matched multi-omic kidney atlas (RNA and chromatin accessibility) derived from the same samples ^10^. To mitigate the inherent sparsity of single-cell methylation data, we constructed metacells by aggregating profiles from the 50 nearest neighbors, yielding an approximate 1X genomic coverage per metacell. Integrating these metacells with multimodal transcriptome and chromatin accessibility data allowed for the distinct characterization of healthy and altered states within tubular cell populations.

### Identification of orthologous genomic regions between species

Putative orthologous genomic regions between the human (hg38) and mouse (mm39) genomes were identified using a reciprocal liftover approach incorporating synteny information. The hg38 genome was segmented into non-overlapping 5 kbp bins.

Mapping was performed using the UCSC liftOver tool with reciprocal best alignment chain files (hg38ToMm39.over.chain.gz, mm39ToHg38.over.chain.gz) and a minimum mapping fraction (-minMatch) of 0.7. Initial hg38 bins were mapped to mm39 and the resulting mm39 regions were subsequently mapped back to hg38. Candidate region pairs were filtered, requiring at least 50% overlap between the original hg38 bin and it’s corresponding reciprocally mapped hg38 region. An additional synteny filter was applied using pre-defined syntenic blocks derived from processed UCSC alignment net data (hg38_mm39_synteny.tsv).

### Cross-species data integration and visualization

Human and mouse meta-cell methylation data were integrated using Scanpy (v1.1) ^45^ and batch-corrected with Harmony (harmonypy v0.0.9) ^21^. After initial filtering to remove orthologous regions with over 90% missing values, missing methylation values were imputed using k-nearest neighbors (KNNImputer ^46^, n=5). The datasets were concatenated, keeping only common orthologous regions between species. Highly variable features (n = 10,000) were selected, scaled, and dimensionality reduced by PCA (50 components). Harmony correction was applied to PCA embeddings to correct batch effects across species. Leiden clustering and UMAP embeddings were computed using the corrected neighborhood graph.

### Differential region analysis and consensus reference panel across species

Leiden clusters resulting from cross-species analysis were mapped to consensus cell types, and regions significantly differentially methylated between each target cell type and the rest were identified separately within each species (Wilcoxon rank-sum test). Regions included in the final conserved reference panel using stringent cross-species criteria: (1) average methylation difference between target cell type and the rest exceeding 0.3 in both species, and (2) methylation differences greater than 0.2 between the target and at least 80% of individual other cell types in both human and mouse.

### Identification of cell-type-specific regulatory elements

To identify putative regulatory elements, we utilized the MethylSeekR R package (v1.5)^47^ to segment the genome based on DNA methylation patterns. Given the sparsity of single-cell data, we first generated pseudo-bulk methylomes for each annotated cell type by pooling reads from all nuclei assigned to that cluster. The algorithm segments the genome into UMRs, characterized by unmethylated CpG islands typically corresponding to promoters (<10% methylation), and LMRs, characterized by CpG-poor regions with intermediate methylation (10–50%) typically corresponding to distal regulatory elements or enhancers. Regions located in undefined genomic sequences (N-masked) or on sex chromosomes were excluded from analysis. To identify cell-type-specific regulatory elements, we performed a comparative overlap analysis across all kidney lineages. A UMR or LMR was defined as “cell-type-specific” if it was identified as a segmented region in the target cell type but was not classified as a UMR or LMR (i.e., remained hypermethylated) in any other cell type. Regions identified as UMRs or LMRs in all annotated cell types were classified as “shared”.

### Differential methylation analysis

Cell type-specific DMRs were identified using a two-step approach. First, candidate DMRs were called with the find_markers function from wgbstools (v0.2.2) ^48^, using parameters requiring a minimum length of 50 bp and at least four CpGs per region. Regions were filtered to retain loci with a minimum sequencing coverage of 3X in each cell type. Cell type-specific regions were defined by an FDR < 0.05, a methylation difference of ≥15% against each distinct cell type, and a ≥20% difference relative to the average of all other cell types. Differential methylation between healthy-state and altered-state epithelial cells was assessed using DSS ^49^. DSS accounts for variability in sequencing depth and biological replicates, providing robust detection of statistically significant DMRs. Regions with an FDR < 0.05 were considered significantly differentially methylated.

### Cell type deconvolution

Cell type proportions were estimated from bulk DNA methylation data using UXM ^19^. Reference cell type markers for UXM were selected from the top 250 cell type-specific marker regions identified from the differential methylation analysis described above. To assess robustness and avoid sample-driven bias in marker selection, we performed a leave-one-out cross-validation (LOOCV). For each iteration, one sample was excluded, cell type markers were re-identified from the remaining samples, and UXM-based cell type proportion predictions were evaluated on the excluded sample. This validation ensured the accuracy and generalizability of our cell type proportion estimates.

### Epigenetic clock analysis

To assess cellular aging, we applied established DNA methylation clocks. To enhance robustness and mitigate sparsity, we aggregated methylation data by cell type for each donor prior to analysis. Specifically, we utilized the principal component (PC)-based implementation of Horvath’s clock, which is retrained to minimize noise inherent to individual CpG measurements ^25^. Given that cell type proportions vary significantly (for example, podocytes representing approximately 2% and proximal tubules comprising 25 to 35% of the total kidney cell population) we evaluated whether DNA methylation clock accuracy was influenced by sequencing depth. For benchmarking purposes, we generated pseudobulk methylome datasets at the sample level and systematically downsampled sequencing coverage to depths of 30X, 20X, 10X, 5X, and 3X. Each downsampling was repeated ten times per coverage level. Subsequently, we calculated DNA methylation age for each dataset and compared these ages with the chronological ages of the donors. Epigenetic age acceleration metrics were also computed as the residuals obtained by regressing DNA methylation age on chronological age.

### Identification of methylation haplotype blocks

Methylation haplotype blocks were identified following the approach described by Guo et al. (2017) ^50^. Briefly, haplotype blocks were defined based on co-methylation between adjacent CpG sites using the linkage disequilibrium metric r2, which measures the consistency of methylation status between pairs of CpGs. Genomic regions containing at least four adjacent CpG sites with pairwise r2 values greater than 0.5 were designated as methylation haplotype blocks. Within each block, we calculated methylation entropy (reflecting variability in methylation states) and the proportion of discordant reads (the fraction of sequencing reads exhibiting inconsistent methylation) using the mHapTk toolkit ^51^. Blocks covered by fewer than five sequencing reads were excluded from downstream analyses to ensure robust quantification. To allow unbiased comparison between healthy and altered cell states, we matched the number of analyzed cells across conditions within each sample, thereby minimizing potential biases arising from differences in sequencing depth.

### Xenium data generation and processing

Spatial transcriptomics data was generated using Xenium In Situ platform (10x Genomics) with the Xenium Prime 5K gene panel. Formalin Fixed Paraffin-Embedded (FFPE) tissue sections (5µm thick) were prepared, hybridized, and imaged according to the manufacturer’s instructions. Raw data was processed using Xenium Analyzer software (v3.0.2.0), and analysis pipeline (v3.0.0.15) to generate transcript coordinate lists and cell segmentation boundaries.

### Xenium data analysis

Following data acquisition, quality control filtering was performed to remove cells with fewer than 5 total transcript counts and genes not expressed in any cell. For cell type annotation, we performed integrative analyses using the Human Kidney Atlas version 2 (HKAv2) snRNA-seq reference ^10^ available from kpmp.org (https://doi.org/10.48698/16dd-vj20). Xenium and single nucleus RNA-seq counts were merged using Seurat (v5.2.1) and integration performed using a sketch subset (50,000 cells/nuclei) using the reciprocal PCA (rPCA) strategy. Clustering was then performed on integrated rPCA embeddings using Pagoda2 (v1.0.12, github.com/hms-dbmi/pagoda2) and clusters projected to the full data set using the ProjectIntegration and ProjectData functions in Seurat. Clusters were initially annotated based upon snRNA-seq labels within each cluster and corresponding cell type marker gene expression profiles. Subgroups of broad cell types (set1: POD, PEC, PT, DTL, ATL; set 2: TAL, DCT, CNT, PC, IC, PapE; set 3: EC, VSM/P, FIB, Ad, Lymphoid, Myeloid, NEU) were then clustered independently using the rPCA and Pagoda2 clustering strategy. Final cell type annotations were assigned to these clusters based on dominant snRNA-seq labels, marker gene profiles and their spatial localization within the tissues.

To identify niches associated with spatially adjacent cell types, we used the BuildNicheAssay function in seurat. For this, cell type co-occurrences within each cell’s neighborhood (neighbors.k = 25) were counted and k-means clustering performed for a specified number of niches (niches.k = 10). Niches and individual cell types were visualized using the ImageDimPlot function in seurat. Identities of niches were assigned based on their composition of cell types, with cell type proportions visualized across all niches using the ggplot2 package (version 3.5.1). Spatial mapping of gene sets was performed using the DecoupleR package (version 2.1.1). For this, spatially weighted gene expression values were used to either directly visualize select gene expression, or to compute gene set enrichments using a univariate linear model (ulm). For transcription factor (TF) activities, TF target gene weight tables were obtained from a previous analysis of the PT-S1/S2 repair trajectory ^10^ using scMega ^52^. Pathway enrichments were performed for Hallmark gene sets (MSigDB). Spatial visualization of TF targets or Hallmark pathways was performed using scanpy (version 1.11.5) for only early repair (8), failed repair (6) and healthy PT (9) niches. Spatial visualization of PT/aPT DMR-associated gene sets and select individual genes (PCK1, PLAU) were for all cells.

Waterfall plots for Gene Set Enrichment Analyses (GSEA) were performed in Seurat (Version 5.3.0) using the SeuratExtend (version 1.2.5) GeneSetAnalysis and WaterfallPlot functions.

### Single-cell methyl-Hi-C library preparation

In parallel to sciMETv2, we applied scMethyl-Hi-C sequencing to a subset of human kidney nuclei to capture both methylation and chromatin contacts. The protocol began with an in situ Hi-C step: nuclei were crosslinked with formaldehyde, permeabilized with SDS, and digested with DpnII. The resulting restriction fragment ends were filled with biotinylated nucleotides and proximity-ligated in situ to join interacting DNA fragments. After ligation, nuclei were stained with DRAQ7 and sorted into 96-well plates (one nucleus per well). Bisulfite conversion was performed directly on the sorted nuclei. The resulting single-stranded DNA was converted into sequencing libraries using a random priming approach: high-concentration Klenow fragments were used to incorporate an internal indexed P5 adapter, followed by the ligation of P7 adapters to the 3’ ends using an Adaptase module. The libraries were PCR amplified with dual-indexed primers and pooled for sequencing.

### Single-cell methyl-Hi-C data analysis

Bisulfite-converted Hi-C sequencing data were aligned to the hg38 reference genome using Bismark ^42^, and ligation junctions were subsequently identified to reconstruct chromatin contacts. As a result, each single nucleus yielded two primary datasets: (1) a genome-wide set of CpG methylation calls, and (2) a sparse Hi-C contact map. For cross-modality integration and cell-type annotation between sciMETv2 and methyl-Hi-C data, we generated sequencing data from matched samples using both modalities. We performed data integration using the run_harmony function from the AllCools toolkit ^53^. Specifically, 5000 highly variable genomic bins (25 kbp each) were selected as integration features. Principal Component Analysis (PCA) was conducted followed by Harmony-based correction to align modalities and accurately annotate cell types across datasets.

### Chromatin compartment analysis

To analyze higher-order chromatin structure, we identified A/B compartments using dcHiC (v2.1) ^54^. First, we generated cell-type-specific pseudo-bulk Hi-C contact maps by aggregating single-cell contact files based on the cell type annotations derived from the DNA methylome analysis. Aggregated contact matrices were constructed at 250 kb resolution. We applied dcHiC to these pseudo-bulk matrices to call chromatin compartments. Briefly, for each chromosome, the contact matrix was normalized, and principal component analysis (PCA) was performed on the correlation matrix to extract the first eigenvector (PC1). PC1 values were used to assign genomic bins to compartment A (active, positive PC1) or compartment B (inactive, negative PC1). The signs of the PC1 values were phased using gene density, such that gene-rich regions were associated with positive PC1 values. To define cell-type-specific and conserved compartments, we compared the compartment scores (PC1) of the target cell type against a reference background constructed from the average of all other kidney cell types. Conserved compartments were defined as bins maintaining the same compartment state across the cell types.

## Data availability

The single-nucleus DNA methylation data (h5ad format), processed cell-type-level methylation profiles (BigWig format), and processed 10X Multiome data generated in this study can be access through the HuBMAP publication page (accession id: HBM233.KJSR.676). The raw sequencing data will be available for download from the database of Genotypes and Phenotypes (dbGaP) (accession id: phs002249) upon acceptance of peer-reviewed manuscript. Publicly available datasets analyzed in this study were obtained from the following sources: human whole-genome bisulfite sequencing (WGBS) data from 205 samples across 39 human cell types were downloaded from GEO under accession number GSE186458; WGBS data from laser-microdissected tubulointerstitium and glomeruli were downloaded from the Kidney Precision Medicine Project (KPMP) atlas (https://atlas.kpmp.org) under DOI 10.48698/hhe6-yv15; human single nucleus 10X Multiome data associated with the HKA was downloaded from KPMP under DOI 10-48698-16dd-vj20; and mouse ischemia-reperfusion injury (IRI) single-nucleus 10X Multiome data were downloaded from GEO under accession number GSE209610.

## Code Availability

The custom scripts and pipeline codes in this study are available on Github at (https://github.com/hrrsjeong/sckidney_methylome_processing; https://github.com/hrrsjeong/sc-kidney-methylome-figures-v0.1)

## Acknowledgements

We thank the WashU Kidney Translational Research Center (KTRC) in part for supporting regulatory approvals, tissue procurement and processing, particularly Amanda Knoten, Kristy Conlon and Amy McMurray. We thank Madhurima Kaushal for data processing. We are grateful for the NIH Common Fund supported Human Biomolecular Atlas Program (HuBMAP) grant U54DK134301. The Kidney Precision Medicine Project (KPMP) is supported by the National Institute of Diabetes and Digestive and Kidney Diseases (NIDDK) through the following grants: U01DK133081, U01DK133091, U01DK133092, U01DK133093, U01DK133095, U01DK133097, U01DK114866, U01DK114908, U01DK133090, U01DK133113, U01DK133766, U01DK133768, U01DK114907, U01DK114920, U01DK114923, U01DK114933, U24DK114886, UH3DK114926, UH3DK114861, UH3DK114915, and UH3DK114937.

We gratefully acknowledge the essential contributions of our patient participants and the support of the American public through their tax dollars. The content is solely the responsibility of the authors and does not necessarily represent the official views of the National Institutes of Health.

## Author Contributions

H.J., S.J., and K.Z. conceived the project. D.D. and X.L. generated single cell methylome sequencing data. S.J., S.R. generated Xenium prime data. S.J. and K.Z. generated the 10X Multiome data. M.T.E. and D.G. generated the whole genome bisulfite sequencing data. H.J., B.B.L., Q.Y., J.P.G. analyzed sequencing data. B.B.L. analyzed spatial transcriptome data. H.J., B.B.L., K.Z. drafted the manuscript. All authors read and approved the manuscript.

