## Supplemental Figure for "A cross-species single-cell epigenome kidney atlas identifies epithelial cells as a driver of epigenetic aging"

**Supplementary Figure 1. Genome browser view of epithelial cell type marker genes.** Fractional methylation levels of CpG loci across cell types for representative marker genes.

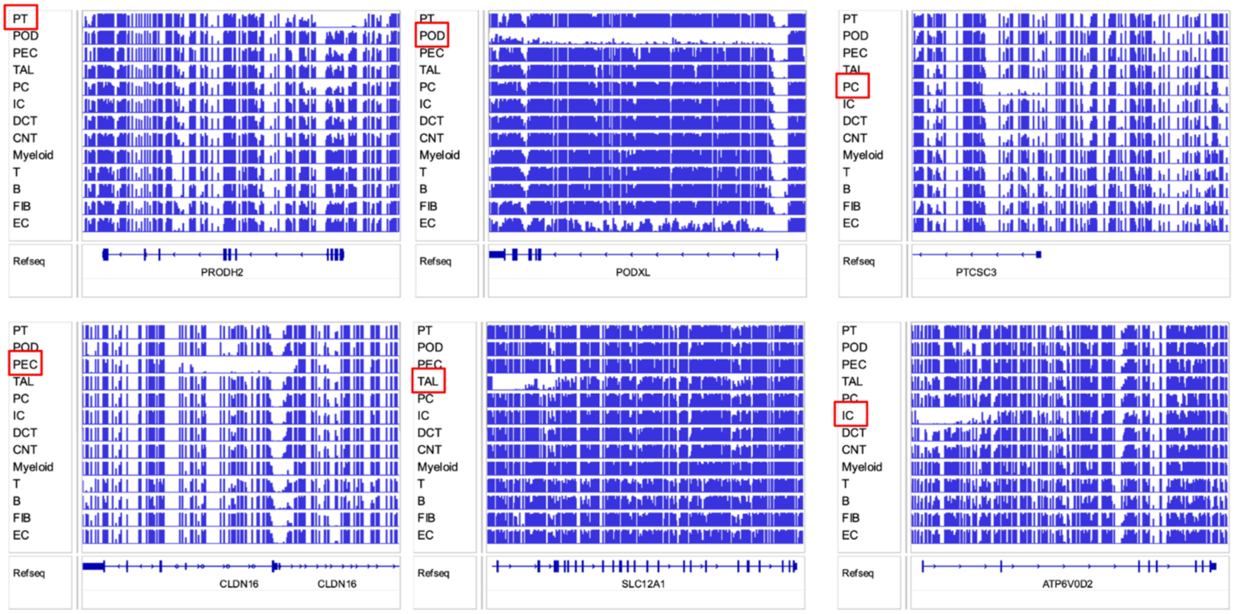

**Supplementary Figure 2.** Principal component analysis of methylation profiles from cis-regulatory regions, comparing with published flow-sorted WGBS data (Loyfer et al., 2023).

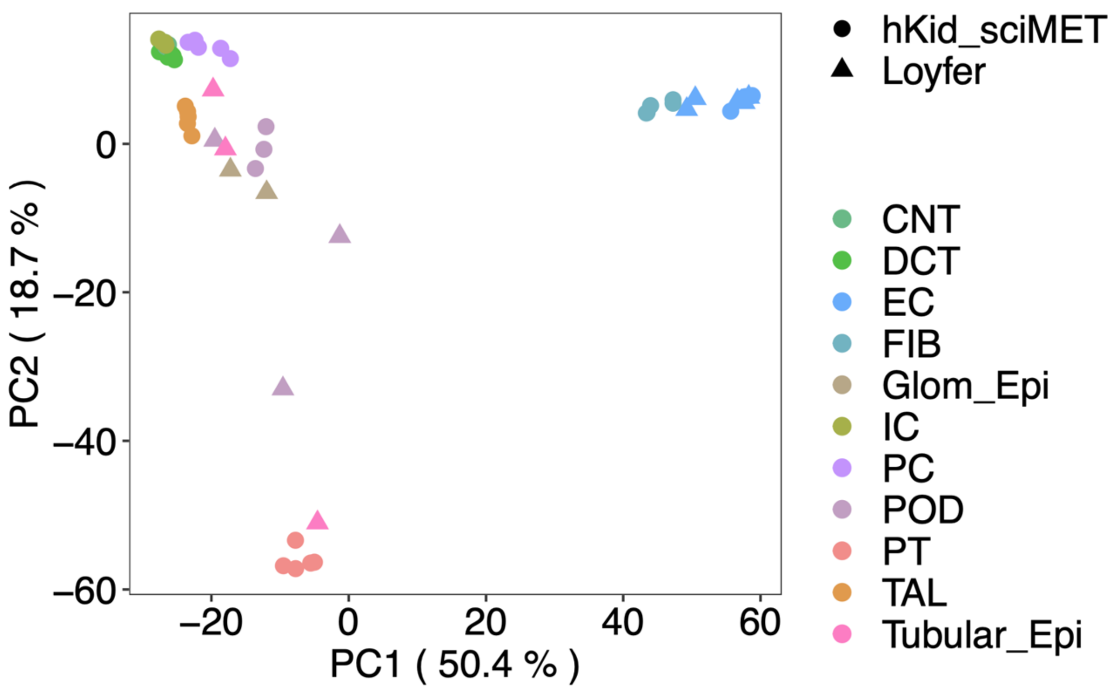

**Supplementary Figure 3.** Box plots of mCG levels for LMRs and UMRs that are unique versus shared across cell types.

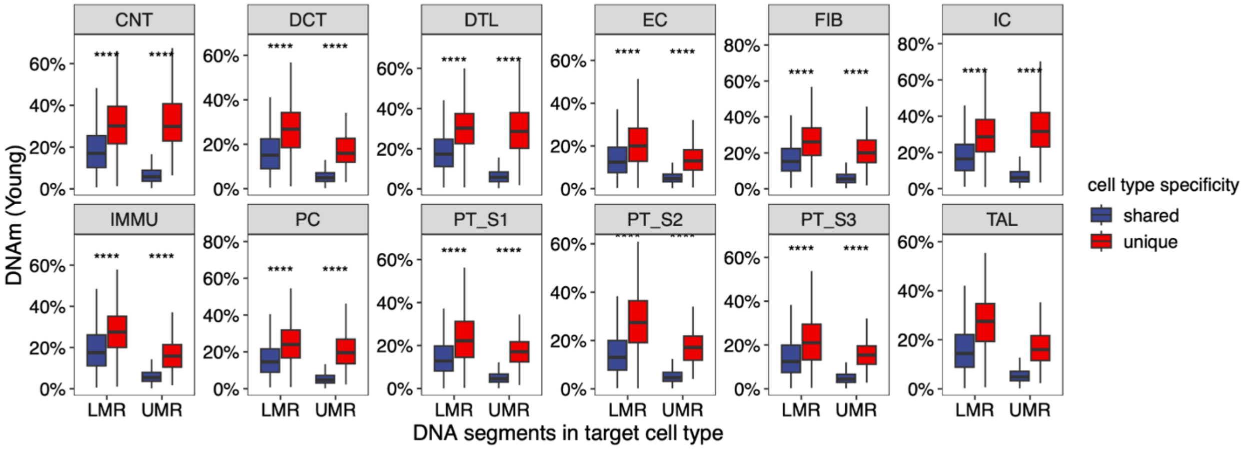

**Supplementary Figure 4.** DNA methylation differences between age groups in lineage-specific versus lineage-shared LMRs and UMRs.

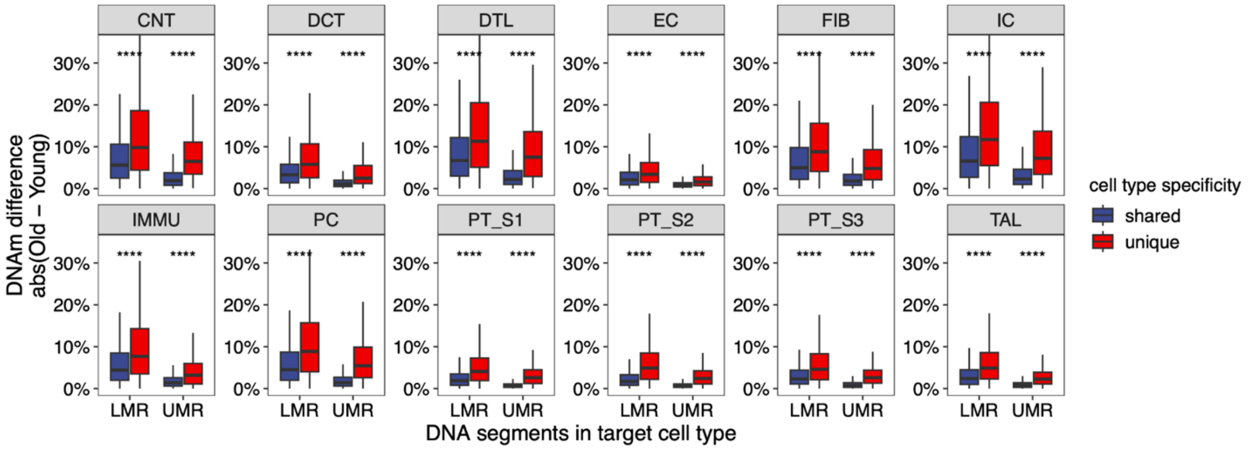

**Supplementary Figure 5.** Correlation between inter-cell-type methylation distance and age-associated methylation differences. For a given target cell type, we first defined its cell-type-specific LMRs/UMRs. Then, for every other cell type, we calculated the average DNA methylation difference between age groups (old vs. young) at these specific loci.

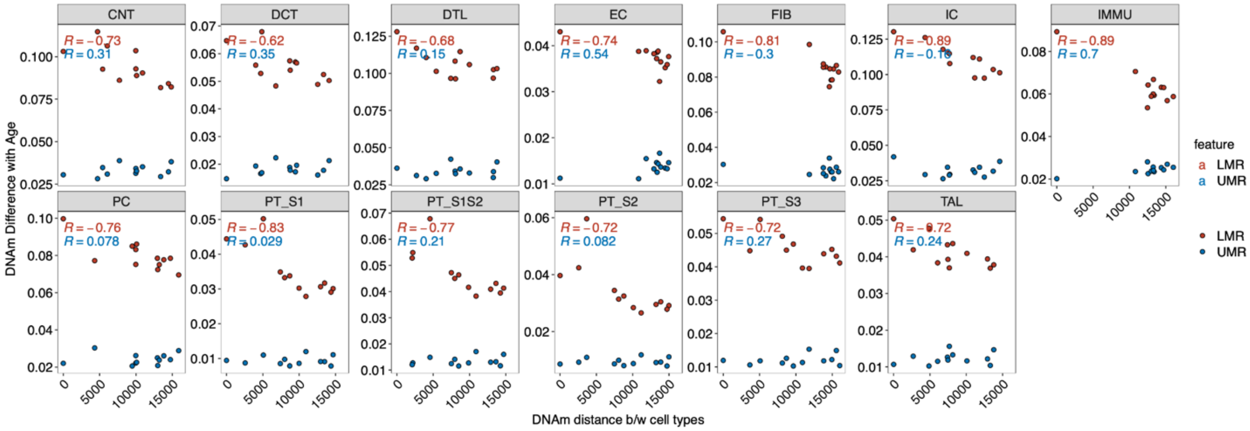

**Supplementary Figure 6.** Proximity test for cross-lineage LMRs. For each target lineage, LMRs belonging to other cell types were ranked by the magnitude of age-associated hypomethylation. The top 10 % ("Aging Hypo") and bottom 10 % ("Least Changed") were compared. Box-plots show the genomic distance from these LMRs to the nearest LMR in the target lineage. Aging-Hypo LMRs from closely related cell types lie significantly closer to the LMRs in the target lineage than those from distantly related cell types, implying that age-associated methylation drift spreads preferentially to regulatory clusters in spatial proximity.

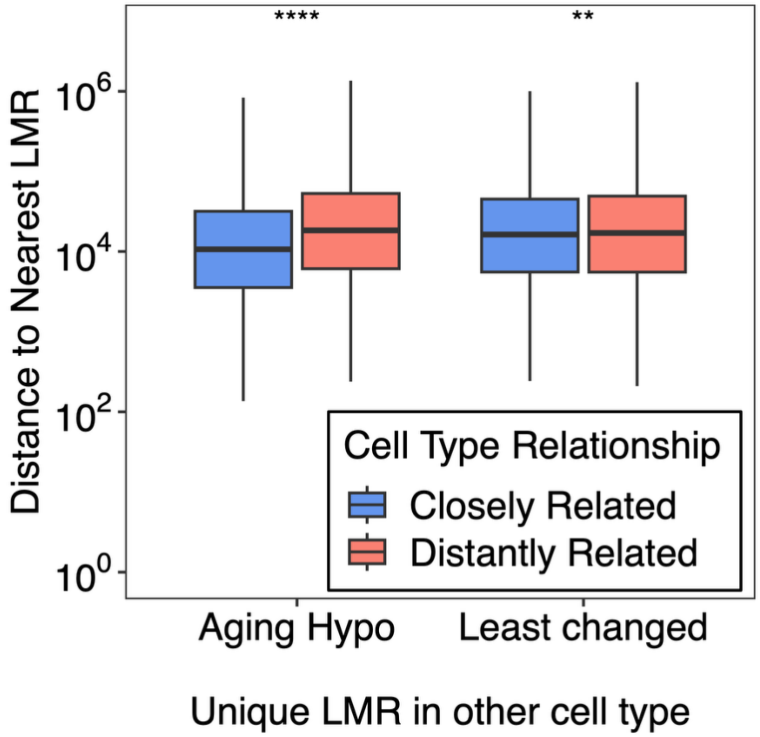

**Supplementary Figure 7.** Robustness of DNA methylation clock analysis at varying sequencing depths. Pseudo-bulk methylomes were generated by downsampling sequencing coverage (3x, 5x, 10x, 20x). This downsampling procedure was repeated 10 times per coverage level, demonstrating that methylation age estimates are stable and accurate across sequencing depths.

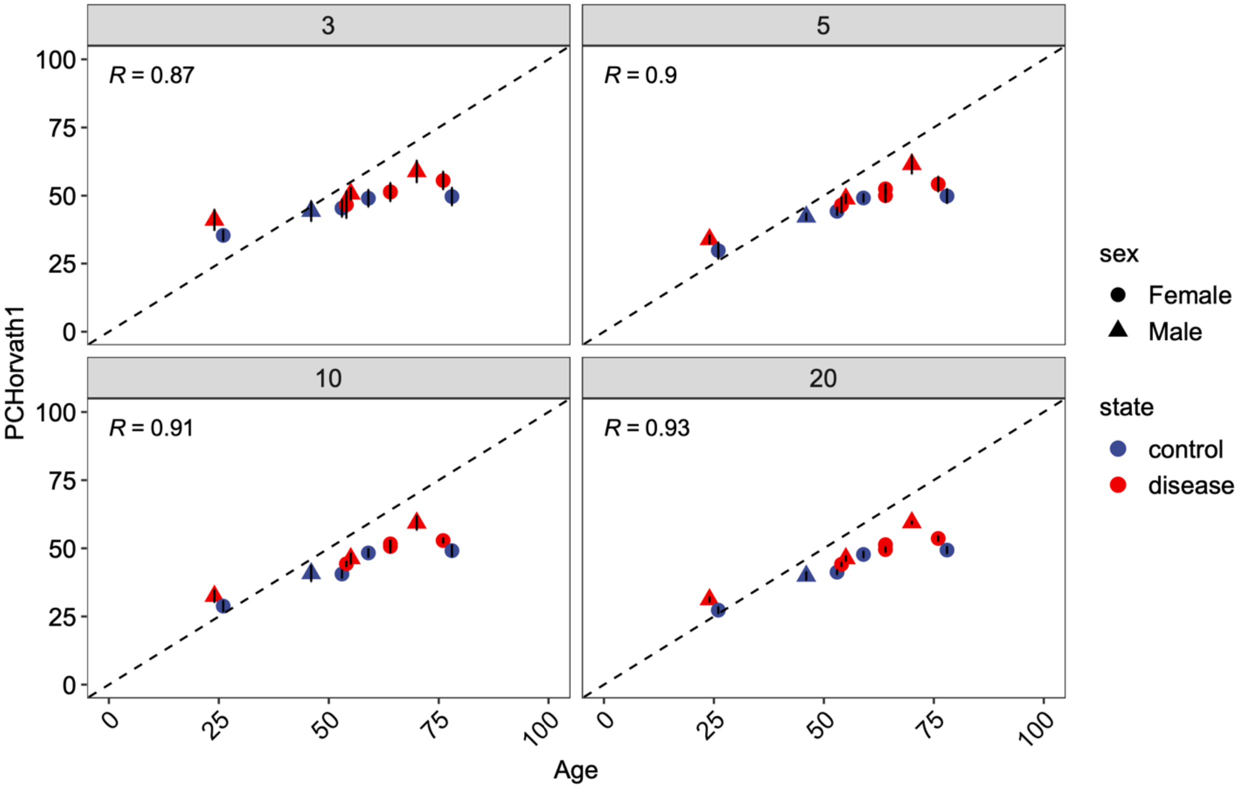

**Supplementary Figure 8.** (a) Cell-type specificity of transcription-factor (TF) motifs in PT ATAC peaks, expressed as (% motif sites in PT-specific peaks / % motif sites in all PT peaks). The ten most PT-enriched motifs are highlighted in red. (b) Comparison of methylation difference between altered and healthy PT at the TF motifs shown in (a). Motifs enriched in PT-specific peaks (red) also show significant methylation gains.

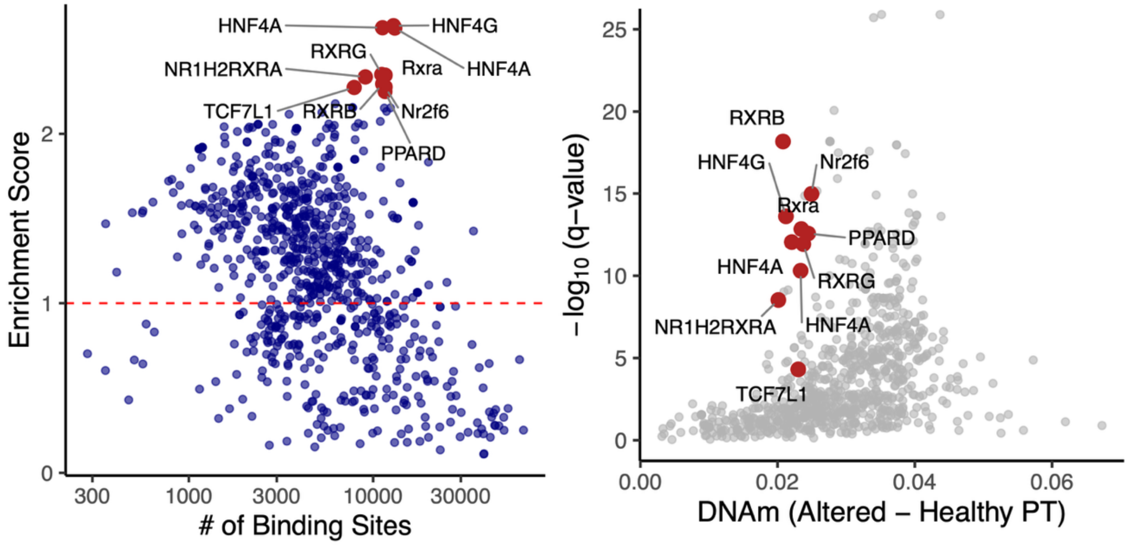

**Supplementary Figure 9.** Transcriptome-wide association with age in PT and TAL. Scatter plots displaying the correlation between gene expression and age derived from 72 healthy control donors. The x-axis represents the correlation coefficient (Spearman's rho), and the y-axis represents the signed significance (signed -log10 P-value). Genes showing significant age-associated upregulation (FDR < 0.1) are colored red, while significantly down-regulated genes are colored blue. Non-significant genes are shown in grey.

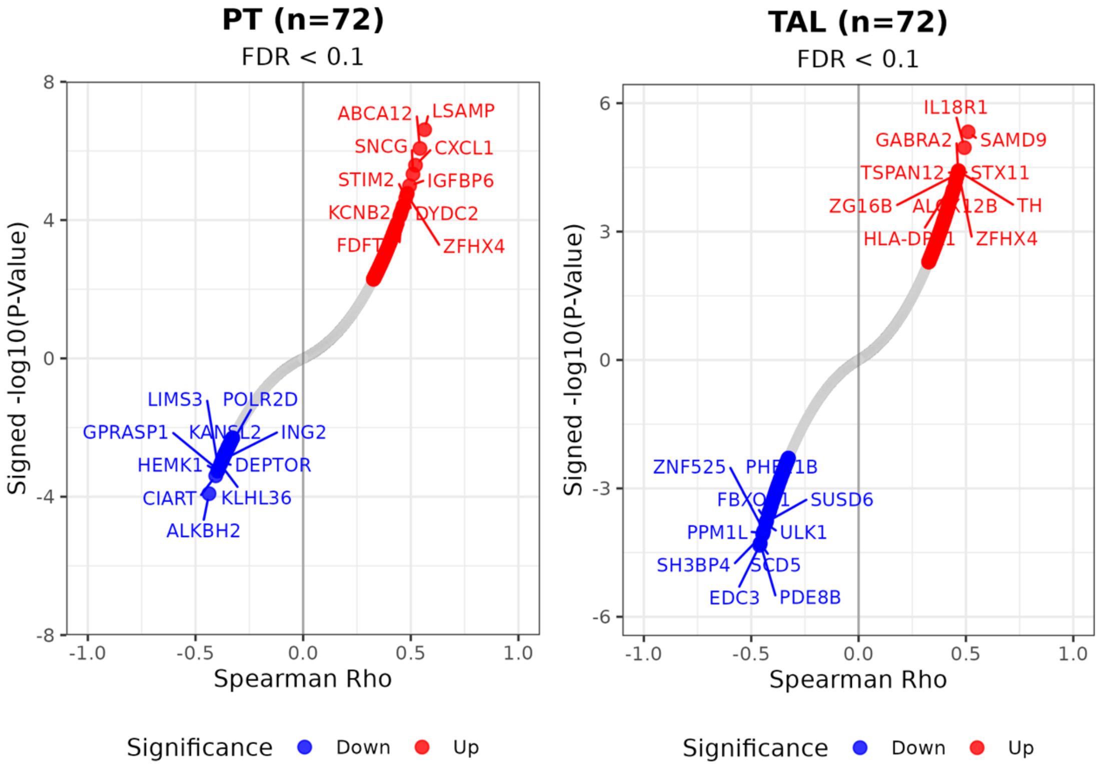

**Supplementary Figure 10.** Top age-upregulated genes in healthy proximal tubule cells. Expression profiles of the top genes positively correlated with age in healthy proximal tubule cells.

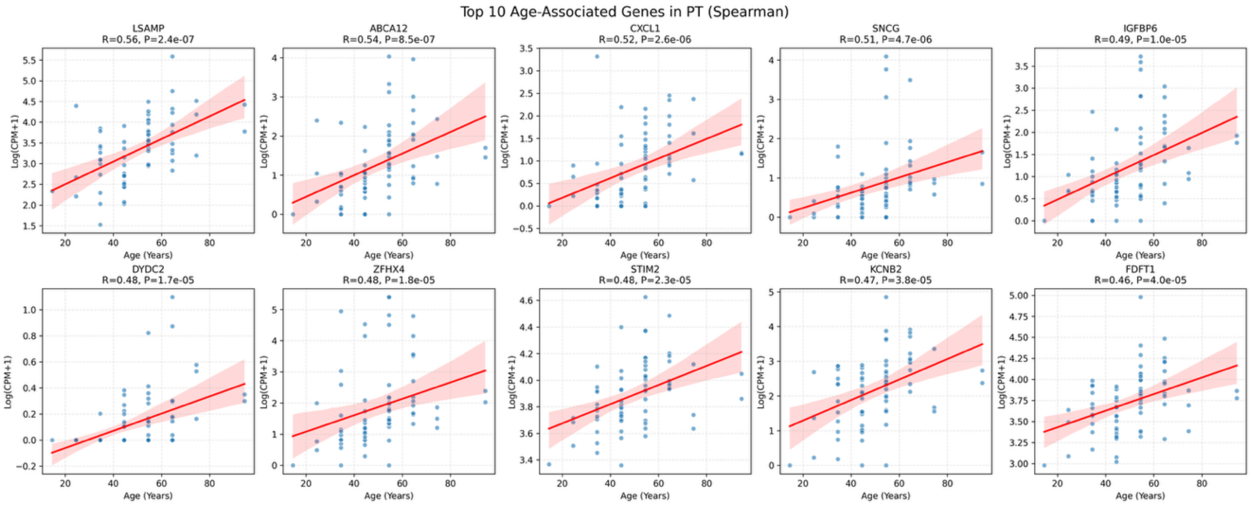

**Supplementary Figure 11. Xenium spatial transcriptomic profiling of human kidney.** Top/middle panels: Cell types and states mapped to healthy reference tissues (HRT) as well as samples associated with acute kidney injury (AKI) and chronic kidney disease (CKD). Epithelial cell types mapped to their expected structures along the corticomedullary axis of the kidney, including the renal corpuscles (POD, PEC), proximal tubules (PT-S1/S2, PT-S3), thin descending and ascending limbs (DTL1, DTL2, DTL3, ATL), thick ascending limbs (M-TAL, C/M-TAL, C-TAL) of the loop-of-henle, distal (DCT) and connecting (CNT) tubules, as well as the collecting ducts (CCD-PC, OMCD-PC, IMCD). Bottom panel: proximal tubule altered cell states associated with early (aPT2) to mid (aPT1) repair states that were enriched in AKI, as well as failed repair (frPT) states that were enriched in CKD.

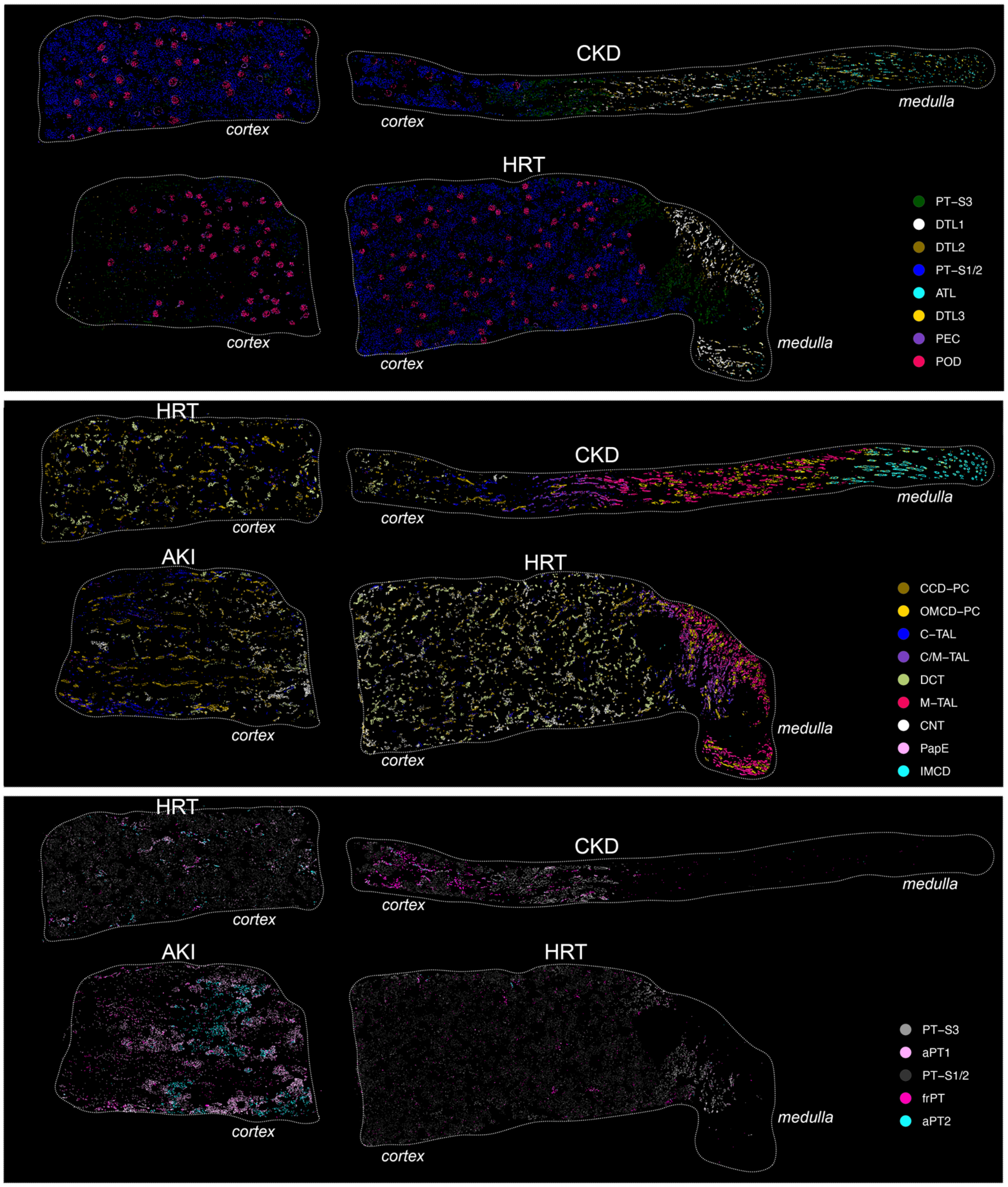

**Supplementary Figure 12. Xenium Niches. a.** Dotplot showing proportions of each cell type across spatial niches (1-10). **b.** Gene set enrichment analyses in proximal tubule-associated niches (9 - healthy; 8 - early repair; 6 - failed repair). Gene set scores plotted for WNT pathway and E2F targets (Hallmark MSigDB), and TF (HNF4A, NFkB2, SOX4) target genes identified from previously defined cell state trajectory gene regulatory networks (GRNs). Arrows indicate early and failed PT repair niches. **c.** Expression scores for PT-aPT DMR-associated gene sets in the AKI sample. Area of early PT repair is highlighted.

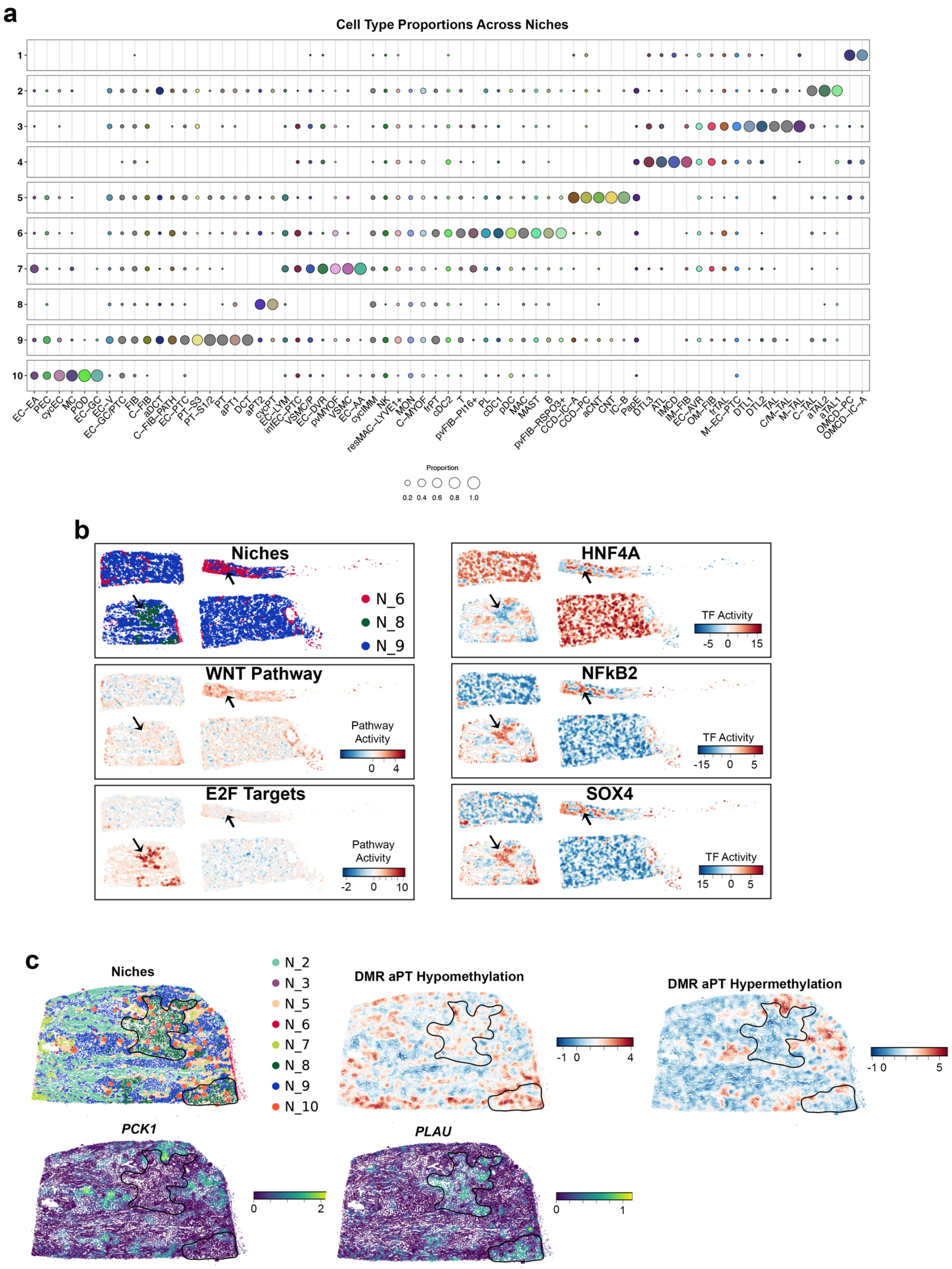
